# Piezo1 and Piezo2 channels in retinal ganglion cells and the impact of Piezo1 stimulation on light-dependent neural activity

**DOI:** 10.1101/2024.06.25.599602

**Authors:** Puttipong Sripinun, Lily P. See, Sergei Nikonov, Venkata Ramana Murthy Chavali, Vrathasha Vrathasha, Jie He, Joan M. O’Brien, Jingsheng Xia, Wennan Lu, Claire H. Mitchell

## Abstract

Piezo channels are associated with neuropathology in diseases like traumatic brain injury and glaucoma, but pathways linking tissue stretch to aberrant neural signaling remain unclear. The present study demonstrates that Piezo1 activation increases action potential frequency in response to light and the spontaneous dark signal from mouse retinal explants. Piezo1 stimulation was sufficient to increase cytoplasmic Ca^2+^ in soma and neurites, while stretch increased spiking activity in current clamp recordings from of isolated retinal ganglion cells (RGCs). Axon-marker beta-tubulin III colocalized with both Piezo1 and Piezo2 protein in the mouse optic nerve head, while RGC nuclear marker BRN3A colocalized with Piezo channels in the soma. Piezo1 was also present on GFAP-positive regions in the optic nerve head and colocalized with glutamine synthetase in the nerve fiber layer, suggesting expression in optic nerve head astrocytes and Müller glia end feet, respectively. Human RGCs from induced pluripotent stem cells also expressed Piezo1 and Piezo2 in soma and axons, while staining patterns in rats resembled those in mice. mRNA message for *Piezo1* was greatest in the RPE/choroid tissue, while *Piezo2* levels were highest in the optic nerve, with both channels also expressed in the retina. Increased expression of *Piezo1* and *Piezo2* occurred both 1 and 10 days after a single stretch *in vivo;* this increase suggests a potential role in rising sensitivity to repeated nerve stretch. In summary, Piezo1 and Piezo2 were detected in the soma and axons of RGCs, and stimulation affected the light-dependent output of RGCs. The rise in RGCs excitability induced by Piezo stimulation may have parallels to the early disease progression in models of glaucoma and other retinal degenerations.

**Highlights:**

- Activation of Piezo1 excites retinal ganglion cells, paralleling the early neurodegenerative progression in glaucoma mouse models and retinal degeneration.
- Piezo1 and Piezo2 were expressed in axons and soma of retinal ganglion cells in mice, rats, and human iPSC-RGCs.
- Functional assays confirmed Piezo1 in soma and neurites of neurons.
- Sustained elevation of *Piezo1* and *Piezo2* occurred after a single transient stretch may enhance damage from repeated traumatic nerve injury.

Graphical abstract
Piezo1 and Piezo2 channels in retinal ganglion cells and the impact of Piezo1 stimulation on light-dependent neural activity. Puttipong Sripinun, Lily P. See, Sergei Nikonov, Venkata Ramana Murthy Chavali, Vrathasha Vrathasha, Jie He, Joan M. O’Brien, Jingsheng Xia, Wennan Lu, Claire H. Mitchell*. Activation of Piezo channels through mechanical or pharmacological stimulation leads to an influx of Ca^2+^ and other cations into RGCs, depolarizing the membrane and increasing the action potential frequency to modulate the visual signal. Created with Biorender.com

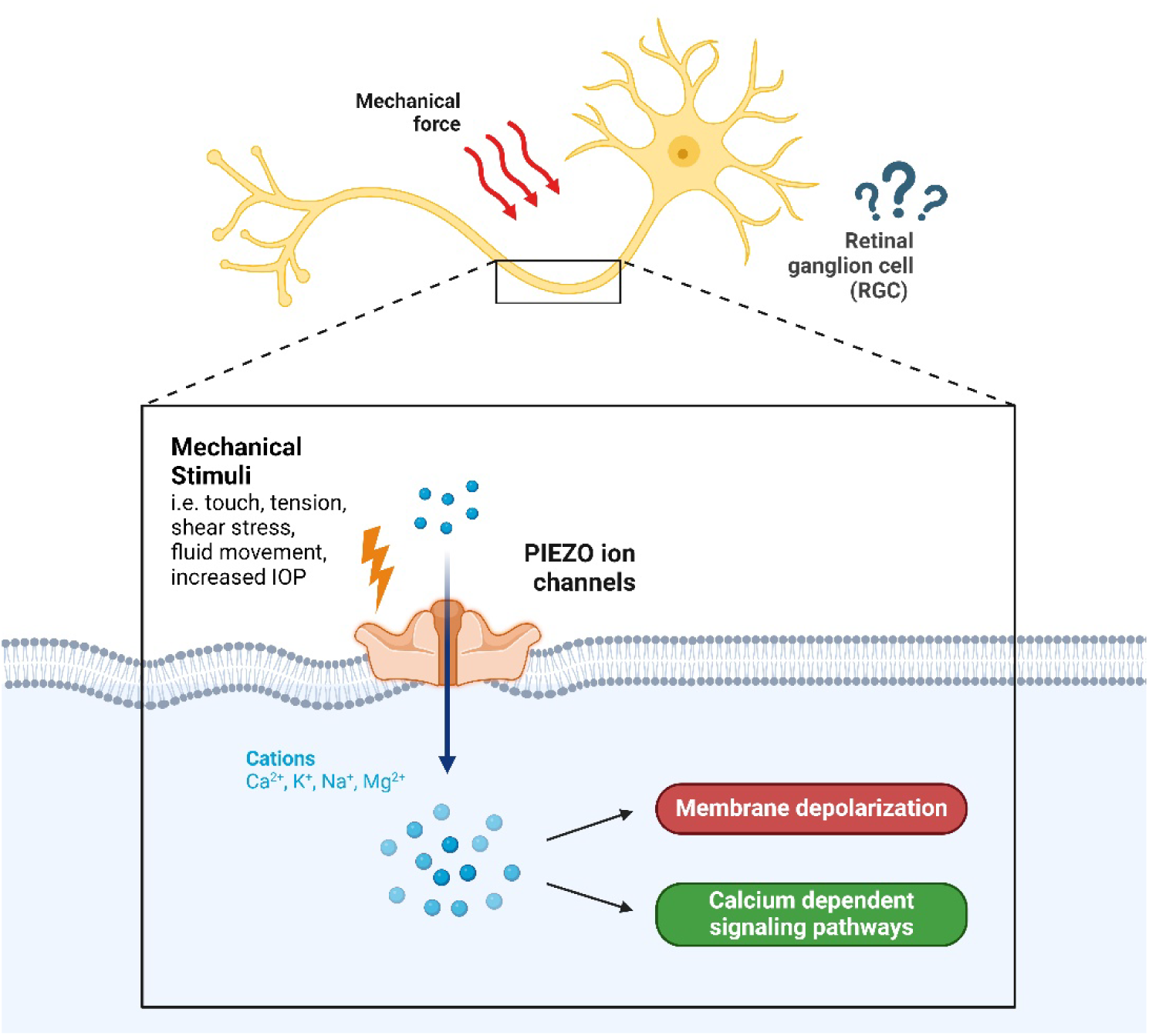

## Introduction

Throughout the body, mechanical stimuli are converted into electrochemical signals through the process of mechanotransduction. Excessive mechanical strain on nervous tissue, whether acute or chronic, can manifest as inflammatory events, loss of neural function, and, in some cases, death of the neuron ^1, 2^. Linking mechanical strain to these responses is thus critical in understanding the signaling pathways that contribute to their pathogenesis ^3, 4^.

Piezo ion channels are pore-forming mechanosensitive cation channels ^5, 6^. They are uncommonly large, consisting of over 2100 amino acids and 24-36 transmembrane regions; the extensive surface area of the channel facilitates the sensing and transduction of mechanical stimuli into downstream signaling events ^7, 8^. The Piezo channels are permeable to cations, where membrane stretch gates the pore opening, leading to influx of Na^+^ and Ca^2+^, among others, causing membrane depolarization under physiological gradients ^9, 10^. There are two Piezo isoforms: Piezo1 was traditionally associated with non-neuronal tissues, and Piezo2 with sensory neurons ^8^. However, synergistic interactions and interdependence between the isoforms suggest these divisions are not binary ^5, 11^. In addition to being mechanosensitive themselves, Piezo channels can mediate the downstream modulation of other mechanosensitive ion channels, making them primary regulators of cellular mechanosensitivity ^12, 13^. The contributions of Piezo channels to mechanotransduction in neural tissues under baseline and stressed conditions remain unclear, with increasing evidence suggesting possible roles of not only Piezo2 but also Piezo1 in neurons ^14–17^.

Retinal ganglion cells (RGCs) are particularly vulnerable to mechanical strain as their path from soma to synapse in the brain requires the unmyelinated axon to transverse the optic nerve head region as they exit the ocular chamber. The mechanical forces driven by the pressure gradients between the cerebrospinal fluid pressure and intraocular pressures (IOP) are greatest here and are exacerbated when IOP rises in glaucoma ^18, 19^. Increased forces across the optic nerve head region remain a primary risk factor for neural degeneration and death of RGCs in glaucoma ^20–23^. While several studies have investigated the presence of Piezo channels in the retinal and optic nerve regions, considerable discrepancies remain ^24–27^. The use of a single species and limited approaches likely added to the confusion. A more rigorous examination of Piezo channels in the neurons and glia of the optic nerve head region is needed, particularly given a recent investigation implicating Piezo1 mutations in some forms of glaucoma ^28^.

The present study uses molecular tools, immunohistochemistry, microelectrode array, and live cell Ca^2+^ imaging to examine the expression of Piezo1 and Piezo2 in the retinal tissue of mice, rats, and human RGCs derived from induced pluripotent stem cells (iPSC-RGCs). Of particular interest is the presence of both Piezo1 and Piezo2 in the axons of RGCs, the response in live RGCs to Piezo1 agonist Yoda1, the rise in *Piezo1* and *Piezo2* expression 10 days after a transient stretch, and increase in the excitability of the RGCs in explant retinal tissue. These findings support the neural expression of both Piezo1 and Piezo2 and suggest that a single stretch can prime neural tissues, increasing their sensitivity for future responses and may potentially alter the visual perception in patients.

## Methods

### Animal Care and Use

Animal experiments were performed in strict accordance with the National Research Council’s “Guide for the Care and Use of Laboratory Animals” and were approved by the University of Pennsylvania Institutional Animal Care and Use Committee (IACUC) protocol #803584. All animals were housed in temperature-controlled rooms under 12:12 light–dark cycles with food and water ad libitum. Long–Evans rats of 12 months (Harlan Laboratories/Envigo, Frederick, MD, United States), and C57BL/6J mice of 3-6 months (Jackson Laboratories, Bar Harbor, ME, United States) of both sexes were utilized.

### iPSC-RGC culture

iPSC-RGC development and growth was performed as described previously with minor modifications ^29^. Human iPSCs were kept on dishes coated with 1:100 diluted growth factor reduced Matrigel (Corning) and maintained in iPSC medium (StemMACs iPS-brew XF, Human, Miltenyi Biotec) with the addition of 20 ng/mL bFGF. Once cells attained 75% confluence; 5 ng/mL stable bFGF was added. iPSCs were kept at 37 °C with 5% O_2_ and 5% CO_2_ until 100% confluent, then transferred overnight to 5% CO_2_ to initiate RPC/RGC induction. On day 0, media was changed to RPC induction media: DMEM/F12 (50:50), 1% P/S, 1% Glut, 1% NEAA, 0.1 mM 2-ME, 2% B27 supplement (w/o vitamin A), 1% N2 supplement. Daily additions of fresh media including 2 μM XAV939, 10 μM SB431542, 100 nM LDN193189, 10 mM nicotinamide, and 10ng/mL IGF1 were performed. Nicotinamide was replaced with 10ng/ml bFGF on days 4-21. RGC induction media was introduced on day 22, composed of DMEM/F12: Neurobasal media, 1:1, 1% P/S, 1% Glut, and 2% B27. In addition, 250ng/ml SHH, 100ng/ml FGF8 and 3uM DAPT were added to media on days 22-23. On day 24, cells were cross-hatched and passaged onto Matrigel coated plates for RGC differentiation with 3µM DAPT, and 100ng/ml Follistatin included in the RGC medium. Cyclopamine 0.5µM was included on day 24 only. RGC maturation was induced on days 27-35 with addition of 3µM DAPT, 10µM Y27632, 400µM cAMP, 5µM forskolin, 40ng/ml BDNF, 5ng/ml NT4 and 10ng/ml CNTF. Cell media was changed daily from days 0-26, and every 3 days from days 27-35. After day 36, RGCs were maintained in RGC induction media with DAPT and Y27632 with media changed 2x/ week. The iPSC-RGCs produced were shown to express markers RPBMS and BRN3A and produce action potentials when depolarized ^29^, supporting their identity as RGCs.

### RGCs in mixed retinal cell culture

RGCs were cultured in mixed retinal cultures as these cells are healthier than those grown under isolated conditions ^30^. To ease the identification of RGCs, cultures were generated from P3-10 *Slc17a6^Cre+^*; *R26R^tdTomato+^*mouse pups. tdTomato-labeled RGCs mice (*Slc17a6^Cre+^*; *R26R^tdTomato+^*), *Slc17a6*-driven Cre (*Slc17a6^Cre+/+^*; strain number 028863) mice were mated with Ai9 mice (*R26R^tdTomato^*; strain number 007909), which express CAG promoter-driven tdTomato expression upon Cre-mediated recombination, thereby removing loxP-flanked STOP cassette allowing for a transcription of red fluorescent tdTomato as described ^31^. Tail DNA was used for genotyping confirmation and analyzed using the primer sequences and PCR protocols following manufacturer guidelines (Jackson Laboratories). To generate mixed retinal cultures, pups from both sexes were euthanized and retinas were promptly dissected in HBSS (Cat# 14175079; Gibco). Retinas were then incubated in digestion media containing papain (16.5U/mL; Cat# LK003176; Worthington) and DNase I (12,500U/mL; Cat# LK003170; Worthington) for 30 mins at 37°C followed by trituration in RGC culture media and centrifugation. Cells were then plated on poly-D-lysine and mouse laminin I-coated 12 mm Nunc glass base dish (Cat# 150680; Thermo Scientific) and cultured for 3-5 days in the RGC culture medium prior to the calcium imaging experiment. RGC culture media was based on the enriched serum-free growth media as described ^32^; RGC growth and survival were substantially improved as compared to approaches used previously in the laboratory ^33^. The media containing a 1:1 mix of base media of DMEM (Cat# 11960044; Gibco) and neurobasal (Cat# 21103049; Gibco), insulin (5 μ g/mL; Cat# I6634; Sigma), sodium pyruvate (1mM; Cat# 11360-070; Sigma), NS21 supplement (Cat# AR008; R&D systems), penicillin/streptomycin (1%, Cat # 15140-12; Gibco), SATO supplement (1X; in house-made) ^34^, L-glutamine (2mM, Cat# 25030-081; Gibco), triiodothyronine (T3, 40 ng/ml, Cat# T6397; Sigma), brain-derived neurotrophic factor (BDNF; 50 ng/mL; Cat# 450–02; Peprotech, Rocky Hill, NJ), ciliary neurotrophic factor (CNTF; 10 ng/mL; Cat# 450–13; Peprotech), and forskolin (4.2ng/mL; Cat# F6886; Sigma Aldrich). Half of the growth medium was changed every 2–3 days.

### Culture of optic nerve head astrocytes

Primary mouse optic nerve head astrocyte cultures were dissected from neonatal mice (PD1-7) of both genders based on a protocol modified from Albalawi et al. ^35^. In brief, optic nerve head tissue was digested in 0.25% trypsin (Invitrogen, Carlsbad, CA) for 30 mins at 37°C, followed by light trituration in culture media; this loosens up the optic nerve tissue to facilitate astrocyte outgrowth. The material was re-suspended in growth media containing DMEM, 10% FBS, 1% penicillin/streptomycin, and 25 ng/mL murine epidermal growth factor (#E4127, Sigma–Aldrich) and plated on 35 mm culture dishes pre-coated with poly-D-lysine and grown at 37°C, 5% CO_2_. Dishes usually approached confluence within 10 days. Astrocytes were generally used at passage 4 or below.

### Microelectrode array (MEA) recordings

MEA recordings were performed following previously described methods ^36, 37^ on C57BL/6J mice dark-adapted for at least 2 hours. Dorsal retinal patches were positioned with the ganglion cell layer facing downward on the square array of 60 electrodes (200 µm interelectrode distance) in the MEA recording chamber (60MEA200/30iR-Ti-gr; Multi Channel Systems) mounted on the 1060i amplifier (Multi Channel Systems, Reutlingen, Germany). Gentle suction was carefully applied to generate negative pressure bellow the sample, improving electrode-to-tissue contact and the signal-to-noise ratio. During the recording, perfusion of oxygenated Ames’ solution (#A1420, Sigma–Aldrich) at 37°C was continuously supplied to the chamber to support the physiological health condition of the retinal tissue. The Piezo1 agonist Yoda1 was added to the perfusate as indicated. Retinal stimulation included series of ten 2-second flashes of 455 nm light (∼60% efficiency in driving mouse rhodopsin compared to 500 nm light) delivered at 0.1 Hz. All flashes in a series had the same intensity, three different intensities in the scotopic range (32, 117, and 377 hv s^-1^ µm^-2^) were used for the stimulation. Data capture was acquired by an NI PCI-6071E DAQ board and custom software developed in LabView (National Instruments, Austin, TX, USA), and later analyzed using MATLAB-based custom coding (MATLAB, Natick, MA, USA).

### Ca^2+^ measurement from RGCs and optic nerve head astrocytes

To determine the physiologic responses from axons and somas, levels of Ca^2+^ in RGCs were determined microscopically, based on approaches described in detail previously ^38^. In brief, mixed retinal cultures were loaded with 5 µM Fura-2 AM (acetoxymethyl ester, #F1221; Thermo Fisher Scientific) with 0.01% pluronic F-127 (#P3000MP; Thermo Fisher Scientific) at 37°C for 45 min. Cells were washed, mounted in a perfusion chamber, and visualized using a ×40 objective on a Nikon Diaphot microscope (Nikon, Melville, NY, USA). Ratiometric measurements were performed by alternating the excitation wavelength from 340 to 380 nm and quantifying emission ≥512 nm with a charge-coupled device camera (All Photon Technologies International, Lawrenceville, NJ, USA); the tdTomato assisted in identifying the RGCs but did not interfere with the Fura-2 signal. Cells were perfused with isotonic solution containing 105 mM NaCl, 5 mM KCl, 6 mM 4-(2-hydroxyethyl)-1-piperazineethanesulfonic acid, 4 mM Na 4-(2-hydroxyethyl)-1-piperazineethanesulfonic acid, 5 mM NaHCO_3_, 60 mM mannitol, 5 mM glucose, and 1.3 mM CaCl_2_; the Piezo1 agonist Yoda1 (Cat# SML1558, Sigma) and gadolinium (Cat# 01128606, Thermo Fisher) was added to the perfusate for the time indicated.

Given the homogeneity of optic nerve head astrocytes grown in isolation, Ca^2+^ was measured using a plate reader. Astrocytes were grown to confluence in 96-well black assay plates with clear bottoms for 4 –7 days (Cat# 353219; Corning) and loaded with Fura-2 AM and 0.02% Pluronic F-127 in culture media for 45 min at 37°C, followed by a 30 min period for de-esterification. Cells were washed and placed in isotonic solution, and changes in cytoplasmic Ca^2+^ levels were determined from the ratio of light excited at 340 and 380 nm (510 nm emission) using a Varioskan™ LUX multimode microplate reader (Thermo Fisher). Yoda1 was introduced using the internal injector system and was not washed off.

### Retinal stretch with IOP elevation in mice

The IOP was elevated in mice using a modified version of the Control Elevation of IOP (CEI) protocol developed by Morrison et al. ^39^ and used previously ^40, 41^. This procedure separates the acute effects of increased IOP from cell death to focus more on the consequences of mechanical strain. In brief, C57BL/6J mice ages 3-6 months were anesthetized and maintained with 1.5% isoflurane throughout the procedure after receiving 5 mg/kg meloxicam. Corneal anesthesia and mydriasis were achieved by administering proparacaine (0.5%) and tropicamide (1%) eye drops, respectively. One eye was cannulated with a 33-gauge needle attached to polyethylene tubing (PE 50; Becton 51 Dickinson) inserted into the anterior chamber, connected to a 20 ml syringe filled with sterile Hank’s Balance Salt Solution. IOP was elevated to 58 mmHg by positioning the reservoir to the appropriate height; this pressure was selected to maintain retinal blood flow and avoid ischemia ^42–44^ Ocular lubricant gel was applied at regular intervals to both eyes to prevent corneal desiccation (GenTeal; Alcon laboratory, Fort Worth, TX). The needle was removed after 4 h and IOP returned to baseline, with 0.5% erythromycin ointment applied to the cornea. The contralateral eye served as a normotensive control. Mice were euthanized 1 or 10 days later. A sham control mice were also conducted where needle insertion without elevation of intraocular pressure was performed in a similar manner as the experimental group.

### PCR and Real-Time qPCR

Murine eyes were enucleated and promptly dissected in HBSS. The anterior portion of the eye was removed, followed by the iris and lens. Three tissues were collected at this stage under the dissecting microscope: the neuronal retina was detached from the eyecup, RPE sheets were gently peeled off from the underlying Bruch’s membrane by using a microsurgical Iris Spatulae, and optic nerve starting from the proximal portion of the sclera. All tissues were stored in TRIzol reagent (Invitrogen) at −80 °C until further processing for RNA. Total RNA was extracted with PureLink™ RNA Mini Kit (Invitrogen) and RNeasy Micro Kit (Qiagen). cDNA was synthesized from 250 and 500 ng of total RNA per reaction using the High Capacity cDNA Reverse Transcription Kit (Applied Biosystems, Waltham, MA) for PCR and RT-PCR experiments, respectively, as described ^40, 45^. PCR amplification of cDNA was performed using AmpliTaq Gold DNA Polymerases with Gold buffer and MgCl_2_ (Thermo Fisher, Carlsbad, CA) following the manufacturer’s instructions. The PCR products were then run on 2% agarose gels and visualized by ethidium bromide staining, along with a 100-bp DNA ladder (Thermo Fisher, Carlsbad, CA). The real-time quantitative polymerase chain reaction (Real-Time qPCR) was performed using SYBR Green and the 7300 RealTimePCR system (Applied Biosystems). All data were analyzed using the delta-delta Ct approach. *Gapdh* was utilized as an endogenous control. The primer sequences and expected product sizes are given in Table 1.

**Table 1.**
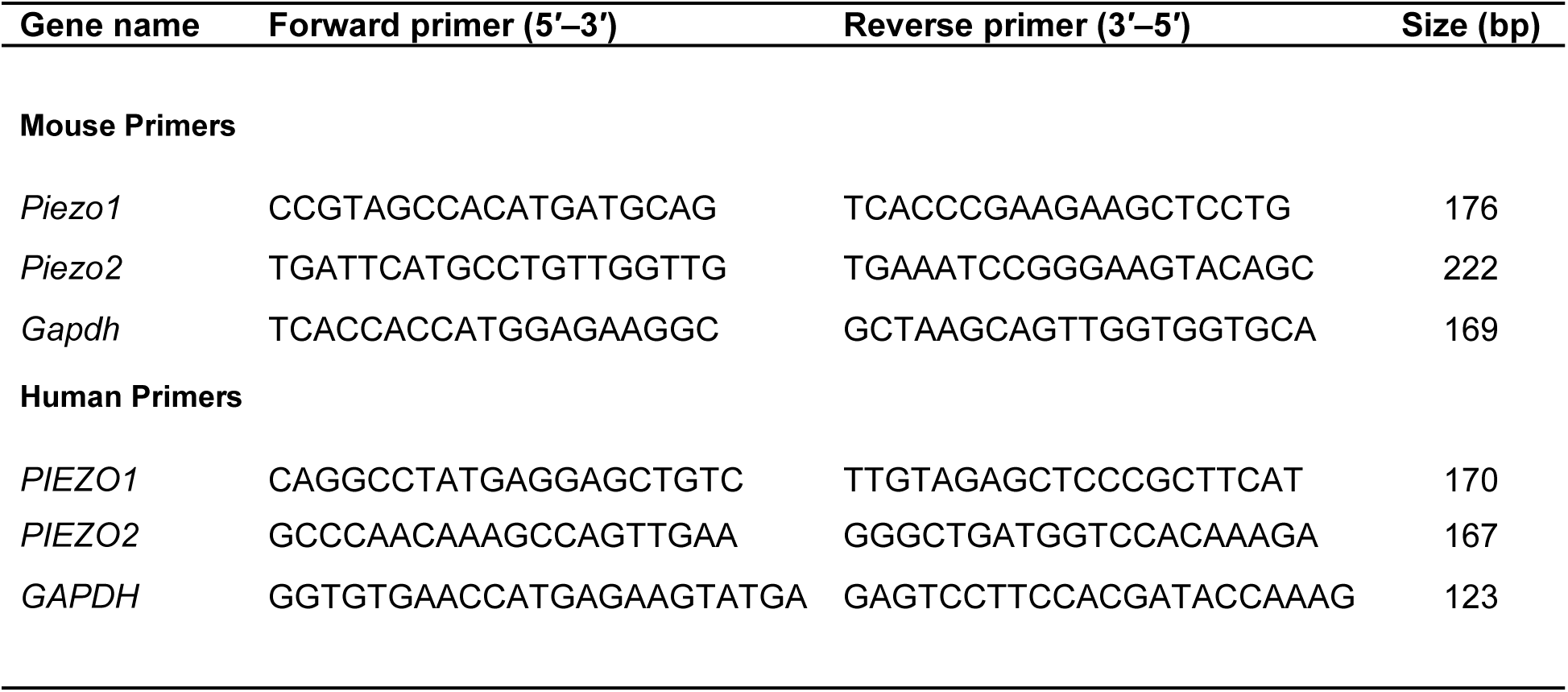
List of primers.

### Immunohistochemistry

Animals were perfused intracardially, and dissected eyes were postfixed with 4% paraformaldehyde. Piezo channels were identified by immunohistochemical staining in 10 µm sections from rodent retina and human iPSC-RGCs. Sections and cells were fixed with 4% paraformaldehyde for 10 minutes, permeabilized with 0.1% Triton X-100 (Sigma-Aldrich) and 20% SuperBlock buffer (ThermoFisher) in 0.1% PBS-T for 10 minutes at 25 °C, then blocked with 10% donkey serum and 20% SuperBlock in PBS-T for 1 hour, followed by primary antibodies overnight at 4°C, and by secondary antibodies for 1 hour at 25 °C. The list of antibodies is provided in Table 2. Autofluorescence quenching was performed on retinal tissues using the TrueVIEW Autofluorescence Quenching Kit (Vector Laboratories, Burlingame, CA) as outlined by the manufacturer. After incubation with DAPI (Sigma-Aldrich) for 10 min, slides were washed and mounted using SlowFade Gold (Thermo Fisher). Imaging was performed using a Nikon E600 epifluorescence and Eclipse confocal Ti2-microscope with NIS Elements Imaging software (Nikon v. 4.60).

**Table 2.** List of antibodies.

| **Antibody** | **Source** | **Identifier** | **Dilution** |
| --- | --- | --- | --- |
| **Primary Antibodies** |  |  |  |
| Rabbit anti-Piezo 1 | Novus Biologicals | Cat# NBP1-78537, RRID:AB_11003149 | 1:100 |
| Rabbit anti-Piezo 1 | ProteinTech | Cat# 15939-1-AP, RRID:AB_2231460 | 1:100 |
| Rabbit anti-Piezo 2 | Novus Biologicals | Cat# NBP1-78624, RRID:AB_11005294 | 1:200 |
| Biotin Mouse anti-Tubulin β3 | Biolegend | Cat# 801212, RRID:AB_2721321 | 1:200 |
| Mouse anti-Brn3a | Milipore | Cat# MAB1585, RRID:AB_94166 | 1:50 |
| Goat anti-Brn3a | Santa Cruz Biotechnology | Cat# sc-31984, RRID:AB_2167511 | 1:200 |
| Rabbit anti-RBPMS | GeneTex | Cat# GTX118619, RRID:AB_10720427 | 1:200 |
| Goat anti-GFAP | Abcam | Cat# ab53554, RRID:AB_880202 | 1:200 |
| Mouse anti-Glutamine synthetase | GeneTex | Cat# GTX630654, RRID:AB_2888230 | 1:200 |
| **Secondary Antibodies** |  |  |  |
| Donkey anti-Rabbit Alexa Fluor 647 | Jackson ImmunoResearch Labs | Cat# 711-605-152, RRID:AB_2492288 | 1:500 |
| Donkey anti-Rabbit Alexa Fluor 568 | Thermo Fisher Scientific | Cat# A-10042, RRID:AB_2534017 | 1:500 |
| Alexa Fluor® 488 Streptavidin | Biolegend | Cat# 405235 | 1:500 |
| Donkey anti-Goat Alexa Fluor 488 | Thermo Fisher Scientific | Cat# A-21206, RRID:AB_2535792 | 1:500 |
| Donkey anti-Mouse Alexa Fluor 488 | Thermo Fisher Scientific | Cat# A-21202, RRID:AB_141607 | 1:500 |

### Data analysis

Statistical analysis was performed using GraphPad Prism software version 10 (GraphPad Software, LLC). Bars represent mean ± standard deviation, with individual data points from each retina, sample, or image depicted as dots, as appropriate. Significant differences between two unrelated groups were assessed by unpaired t-test; paired Student’s t-tests were employed when comparing the eyes of the same mouse. One-way ANOVA followed by Tukey’s tests was used to compare three or more means. Statistical significance showed at p-value > 0.05 = ns, * p-value < 0.05, ** p-value < 0.01, *** p-value < 0.001, and ****p < 0.0001.

## Results

### Piezo1 and Piezo2 messages are expressed in mouse retina, RPE/Choroid, and optic nerve and sustainably upregulated with a transient elevation of IOP in vivo

To provide an overview of Piezo message levels in the adult mouse, initial experiments examined the relative expression of mRNA for *Piezo1* and *Piezo2* in regions of the posterior mouse eye. Traditional PCR demonstrated the presence of *Piezo1* mRNA in the retina, RPE-choroid, and optic nerve regions of C57BL/6 mice. Bands were of the expected size of 176 base pairs, with the most intense band from the RPE-choroid tissue. PCR also showed that *Piezo2* was expressed in the retina, RPE-choroid, and optic nerve regions, with the predicted size of 222 base pairs; expression was greatest in material from the optic nerve and RPE-choroid (Fig. 1A).

**Figure 1.**
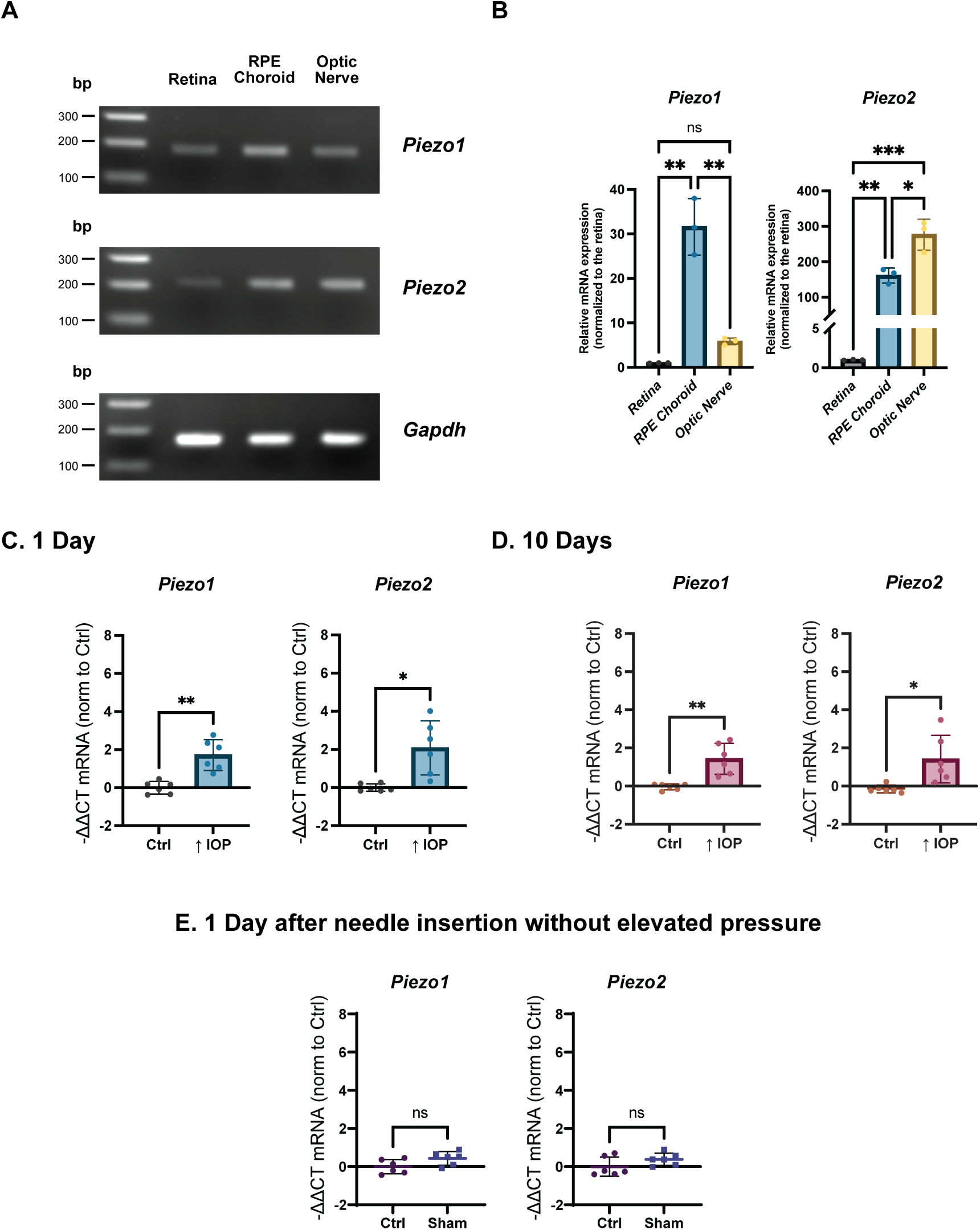
Both mechanosensitive Piezo isoform genes are expressed in the mouse retina, optic nerve and RPE/choroid and were upregulated with transient IOP elevation *in vivo* at both 1 and 10 days. **(A)** Representative 2% agarose gel image of end-point PCR products of mRNA encoding *Piezo1* and *Piezo2*. *Gapdh* served as a loading control. **(B)** Semi-quantitative real-time PCR of *Piezo1* and *Piezo2* expression levels in different tissues in the eye normalized to neural retina. The relative expression of putative mechanosensitive ion channels was in agreement between PCR and semi-quantitative real-time PCR, where both *Piezo1* and *Piezo2* expression levels are expressed in all three types of tissues with greater prominent in RPE choroid and optic nerve tissue when compared to neural retina. (Unpaired t-test; n = 3; * p-value < 0.05, ** p-value < 0.01, *** p-value < 0.001). **(C, D)** Significant upregulation in the expression of both *Piezo1* and *Piezo2* was detected and maintained in recovery time points of 1 and 10 days in retinae exposed to elevated IOP in C57Bl/6J mice. **(E)** No changes were detected with sham needle insertion without elevation of intraocular pressure control group. Dots represent changes in expression from a single mouse, with expression normalized to the average ΔΔCT value of unpressurized contralateral eyes (paired Student’s t-test, n = 6 mice per group).

To further analyze the expression of *Piezo1* and *Piezo2* messages in the posterior eye, we utilized real-time semi-quantitative PCR (qPCR). The relative expression of *Piezo1* mRNA was greatest in material from the RPE-choroid, with levels 30-fold greater than in the retina. Levels of *Piezo1* expression were 5-fold greater in the optic nerve than in the retina. *Piezo2* gene expression was highest in the optic nerve, with expression >200-fold higher than in the retina, while expression of *Piezo2* was 160-fold higher in the RPE-choroid than in the retina (Fig. 1B). The convergence between results from traditional PCR and qPCR regarding the relative expression of Piezo message strongly supports the finding that *Piezo1* and *Piezo2* are expressed in all three types of tissues, with the greatest expression in RPE choroid and optic nerve tissue, respectively, when compared to the neural retina.

Stretch is recognized as a key force impacting optic nerve head and retinal tissue following the elevation of IOP ^46, 47^. To determine whether a single exposure to stretch altered the expression of Piezo channels, a modified version of the Controlled Elevation of IOP (CEI) procedure was used to examine the timescale of Piezo channel upregulation; the CEI procedure can induce RGC damage following a single interval of pressure elevation that resembles that found with more chronic rises ^39^. Mice were sacrificed 1 day or 10 days after a 4-hour hypertensive episode, as a recent investigation suggests inflammatory signals can remain elevated 10 days after a single elevation of IOP ^48^.

A significant upregulation in the expression of both *Piezo1* and *Piezo2* was found in mouse retina stretched with the CEI procedure. *Piezo1* and *Piezo2* expression levels increased 1.7-fold and 2-fold, respectively, 1 day after the return to baseline IOP levels (Fig.1C). Both Piezo isoforms remained significantly elevated 10 days after the event (Fig. 1D). Sham control with the needle inserted in a similar fashion as the experimental group while no elevation of intraocular pressure was used as control where no significant changes can be seen (Fig. 1E). Together, these findings suggest an increase in *Piezo* expressions after a single IOP elevation that could prime the retina, producing increased responses to subsequent increases in pressure.

### Electrophysiological study using ex vivo retinal MEA recording revealed enhanced RGC excitability via Piezo1 activation

Specific agonists that can selectively activate Piezo channels are of great interest and are crucial for examining the roles of Piezo channels in physiological and pathological processes. However, currently, only Piezo1-specific agonists have been identified, while Piezo2 remains an area of ongoing research. To determine the effect of Piezo1 ion channel activation on RGC physiological function, a specific Piezo1 agonist, Yoda1 was used in an established MEA experiment to assess any alteration in spike firing of RGCs from retinal explants. During the course of the experiment, continuous spontaneous firing was notably observable during periods of rest, while substantial activity in the firing of RGCs was elicited at the commencement and conclusion of the light stimulus. With the perfusion of Yoda1 to the system, mean spontaneous firing rates of RGCs significantly increased by 41.16% from the control. This phenomenon continued as Yoda1 remains in the perfusion (Fig. 2A,B) and can be partially alleviated by removing Yoda1 from the system (Fig. 2B).

**Figure 2.**
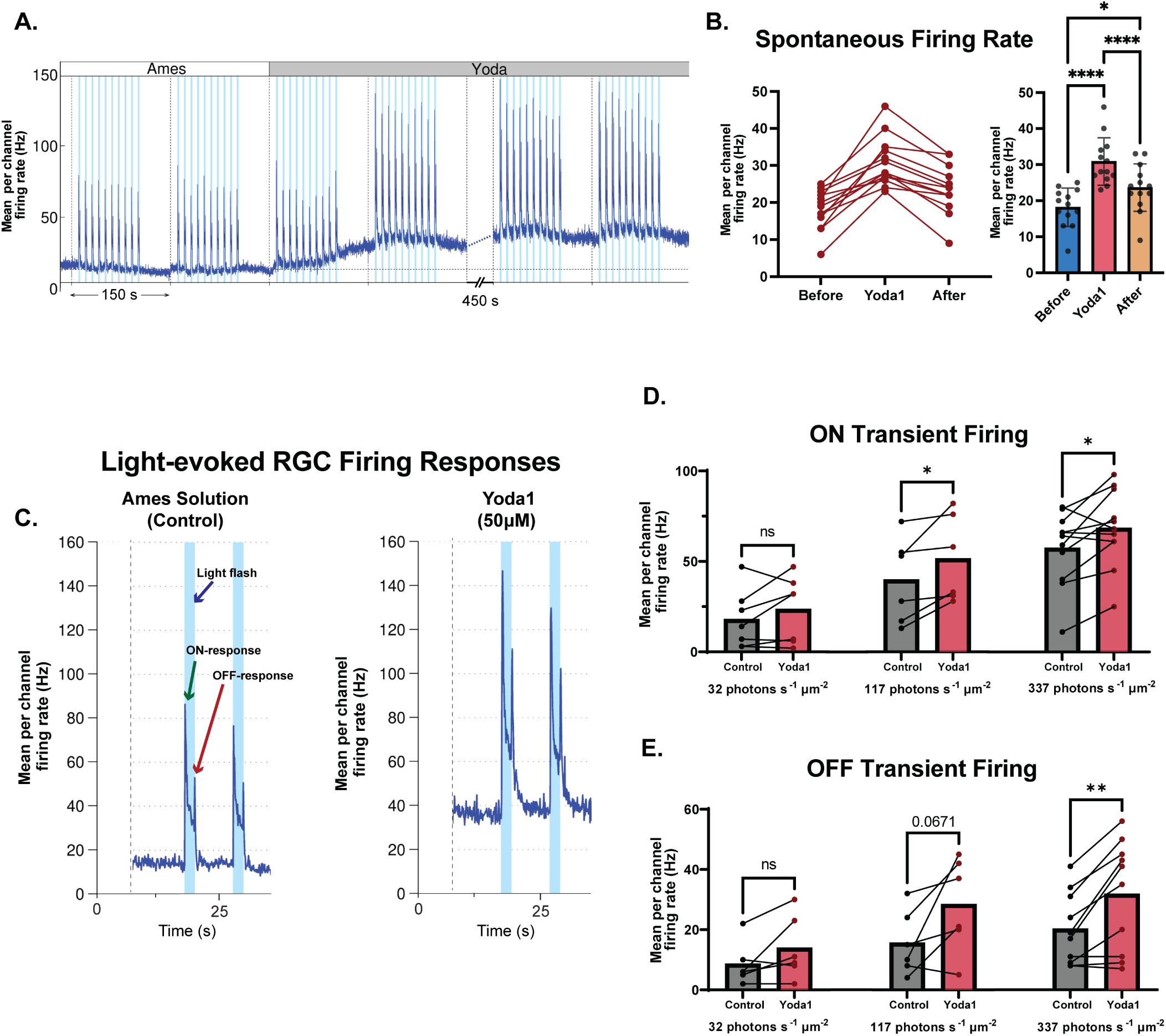
Piezo1 activation altered scotopic RGC responses in retinal explant using MEA recording. **(A)** Representative light-evoked MEA records transitioning from Ames solution to Yoda1, where RGC demonstrated increases in spontaneous firing baseline and enhanced response to 337 photons s^-1^ μm^-2^ flashes of 455 nm light. **(B)** Average spontaneous firing rate of RGCs compared among three-time points. The spontaneous firing rate was increased when challenged by Yoda1 and decreased with the removal of Yoda1. (Repeated measures one-way ANOVA followed by Tukey’s post-hoc test; n = 13 retina). **(C)** Representative light-evoked MEA recordings from control (left) and Yoda1 (right), where RGC demonstrated increases in spontaneous firing baseline and enhanced response to light of both ON- and OFF-responses can be observed. **(D)** On-transient responses from different retinal samples. (Paired t-test; n = 6-11 retina). **(E)** OFF transient responses from different retinal samples (Paired t-test; n = 6-10 retina). Statistical significance showed at * p-value < 0.05, ** p-value < 0.01, and **** p-value < 0.0001.

To further examine the effect of Piezo1 activation on the scotopic condition, a series of ten 2-second flashes at different intensities were used to evoke ON- and OFF-RGC firing activities (Fig. 2C). Analysis revealed a modest but significant increase in light-evoked ON and OFF RGC transient firings (Fig. 2D,E). These findings support the evidence of Piezo1 channel expression in the retina, and its activation can heighten the excitability of RGCs, suggesting high mechanosensitivity within these neuronal cells mediated through Piezo1 channel. Further support comes from patch clamp trials indicating that RGCs display pressure-sensitive currents and increase spiking activity when pressure is applied through a high-speed pressure clamp (Fig. S1, S2). This is consistent with the increased spiking activity found with Yoda1.

### Physiological confirmation of Piezo1 in RGCs and optic nerve head astrocytes

A functional confirmation of Piezo1 in RGCs was determined utilizing Yoda1, a specific activator of Piezo1, with an EC_50_ at mouse Piezo1 of 17.1 µM and no reported activation of Piezo2 at levels under 100 µM ^49^. Measurements were obtained from RGCs in mixed retinal cultures from *Slc17a6^Cre+^*; *R26R^tdTomato+^* mice to aid in identifying RGC soma and neurites; inspection indicated extensive growth of neurites (Fig. 3; A1). RGCs were loaded with the Ca^2+^ indicator Fura-2 as the low excitation wavelengths of 340 and 380 nm did not interfere with the tdTomato signal, while the ratiometric signal was more reliable than single wavelength reporters. Microscopic examination enabled measurement from soma and neurites. Application of 50 µM Yoda1 led to a rapid rise in intracellular Ca^2+^ (Fig. 3; A2). The response was reversible, with levels returning to baseline before Yoda1 was removed. The rise in cytoplasmic calcium observed in both soma and neurites of the RGCs was similar, with no significant difference in magnitude between them (Fig. 3; A3). In addition, a broad-spectrum mechanosensitive cation antagonist, gadolinium (Gd^3+^), was able to inhibit yoda1-induced intracellular calcium elevation (Fig. 3; A4,5).

**Figure 3.**
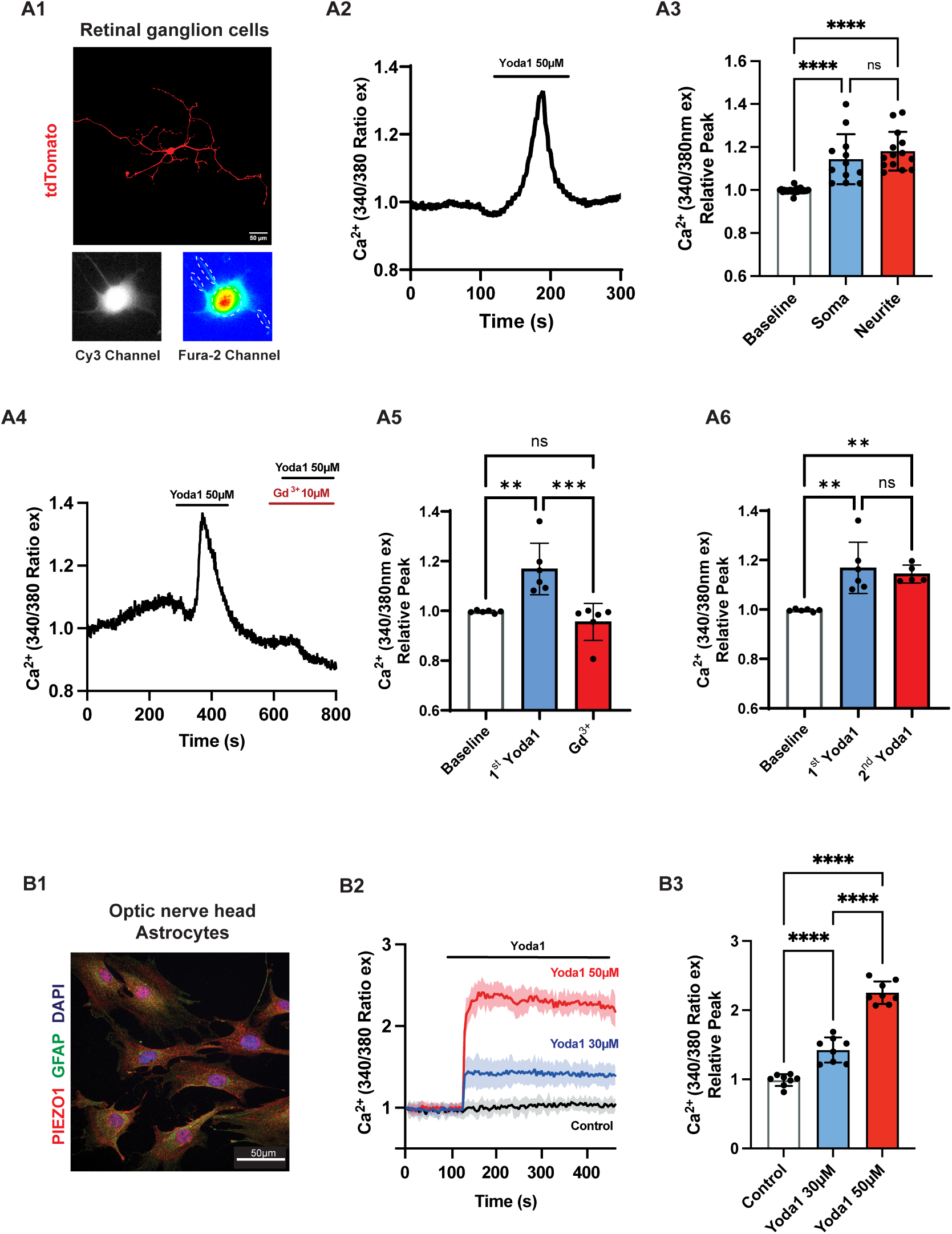
Calcium elevation triggered by specific Piezo1 agonist, Yoda1, in primary mouse RGCs and optic nerve head astrocytes. **(A1)** tdTomato-labeled RGC isolated from *Slc17a6^Cre+^*; *R26R^tdTomato+^* retina. **(A2)** Representative single-cell fura-2 Ca^2+^ imaging trace from a RGC soma in response to 50 μM Yoda1; the duration of drug application is indicated by the horizontal bar. **(A3)** Scatterplot of quantified Fura-2 peak in the response to Yoda1 at soma and neurite as comparison to baseline (Kruskal-Wallis test, n = 12-14; 7 biological replicates). **(A4)** Representative single-cell fura-2 Ca^2+^ imaging trace from a RGC soma in response to 50 μM Yoda1 with or without 3 μM Gd^3+^; the duration of drug application is indicated by the horizontal bar. **(A5)** Scatterplot of quantified Fura-2 peak in the response to Yoda1 as comparison to baseline with or without 3 μM Gd^3+^ (One-way ANOVA tests followed by Tukey’s post-hoc test; n = 6; 2 biological replicates). **(A6)** Scatterplot of quantified Fura-2 peak in the response to repeated Yoda1 exposure (One-way ANOVA tests followed by Tukey’s post-hoc test; n = 5-6; 2 biological replicates). **(B1)** Isolated mouse optic nerve head astrocytes (MONHA) with immunocytochemistry confirmed expression of astrocyte marker, GFAP (green), and Piezo1 (red). **(B2)** Representative Fura-2 Ca^2+^ trace of optic nerve head astrocytes in response to 30 and 50 μM Yoda1 stimulation **(B3)** Scatterplot of quantified Fura-2 relative peak in response to Yoda1 in the astrocytes (One-way ANOVA tests followed by Tukey’s post-hoc test, n = 8; 2 biological replicates).

Further validation of the response to Piezo1 stimulation was examined in mouse optic nerve head astrocytes. The astrocytes stained clearly for both GFAP and Piezo1 (Fig. 3; B1). Staining for Piezo1 was particularly evident on the astrocyte extensions. A similar staining was found for Piezo2 (not shown). Stimulation of the astrocytes with Yoda1 induced a dose-dependent elevation of intracellular Ca^2+^ (Fig. 3; B2,3). The rise in Ca^2+^ induced by Yoda1 in the astrocytes was sustained in the continued presence of Yoda1.

### Detection in RPE cells and validation of antibody specificity

Upon identifying transcripts and functionality of Piezo channels in the retina and considering their potential role in modifying the function of RGCs, we proceeded to investigate the spatial distribution of Piezo channels within the retinal tissues using immunohistochemistry. Although antibodies developed in the past few years have shown improved specificity for Piezo1 and Piezo2 with several genetic modification tool evaluations to support the claims ^50–57^, validation is always needed, so the specificity of the Piezo antibodies used in this study was confirmed. The human retinal pigment epithelial cell line (ARPE-19) was treated with *Piezo1-*targeted siRNA for 48 hours; *Piezo1* siRNAs reduced Piezo1 expression by > 85%. Robust staining for Piezo1 was detected in RPE cells under control conditions but reduced with *Piezo1-*targeted siRNA (Fig. S3B). Quantification of fluorescent intensity confirmed a significant reduction (Fig. S3C). Similar results were found with siRNA targeting *Piezo2*, with a 77% reduction in message (Figure S3D) and decline in immunostaining (Fig. S3E,F). These observations suggest the antibodies used in this study show reasonable specificity, in agreement with previous confirmation reported ^50–57^, while also confirming the detection of message for both *Piezo1* and *Piezo2* in the RPE/choroid tissue above.

### Immunohistochemical identification of Piezo1 in mouse neural retina and optic nerve

The validated antibody was used to examine Piezo1 expression in the adult C57BL/6 mouse retina. Staining for Piezo1 was evident in the nerve fiber layer and cell bodies located in the ganglion cell layer (Fig 4A). Overlap between staining for Piezo1 and glutamine synthase was observed along the internal limiting membrane at the nerve fiber layer (Fig. 4A), suggesting the presence of Piezo1 on the end feet of Müller cells ^58^. Further investigation was proceeded to rule out cell types with positive staining of Piezo1 in the ganglion cell layer.

**Figure 4.**
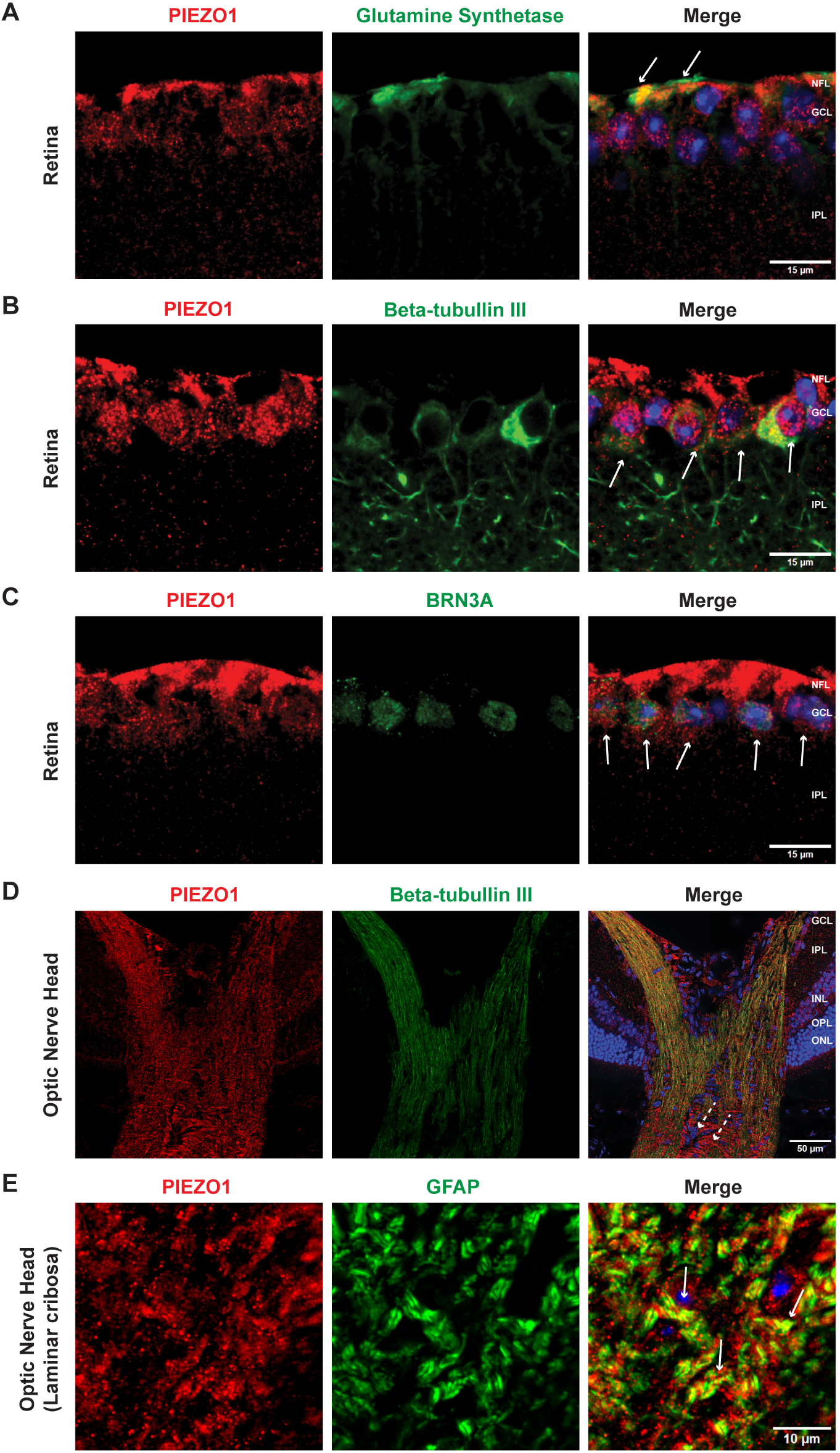
Piezo1 channel immunoreactivity in the mouse retina overlaps with RGC, Müller cells and astrocyte markers: **(A)** Distribution of Piezo1 expression in the internal limiting membrane was highly co-localized with glutamine synthetase, a marker for Müller cells, at the end feet compartments (white arrow). **(B, D)** Robust expression of Piezo1 in the retina in NFL and GCL. Double labeling with anti-Piezo1 and anti-beta-tubulin III antibodies showed positive staining in RGC. **(C)** Representative images of Piezo1 and BRN3A co-staining in the retina demonstrated a shared localization of both proteins in the RGCs’ soma. **(E)** Positive staining of Piezo1 can be seen in the ONH area, where co-staining with GFAP representing astrocytes is evident at the glial lamina in the ONH. Here and throughout, NFL: Nerve fiber layer; GCL: Ganglion cell layer; IPL: Inner plexiform layer; INL: Inner nuclear layer; OPL: Outer plexiform layer; ONL: Outer nuclear layer. Images are representative of the staining found in 3 mice.

The neuronal microtubule marker, beta-tubulin III, was used to identify RGCs ^59, 60^. Positive co-staining between beta-tubulin III and Piezo1 was observed around the RGC cell bodies, though lesser degree of Piezo1 was found in RGC dendrites and axons (Fig. 4B). Close association between Piezo1 and RGC nuclear marker BRN3A ^61^ also suggested expression of Piezo1 in RGC cell bodies (Fig. 4C). Similar immunolabeling was also performed with another validated antibody by other studies where staining in RGC cell bodies can also be seen, supports the premise of Piezo1 expression in RGCs (Fig. S5).

The optic nerve head also displayed staining for both Piezo1 and beta-tubulin III, with colocalization along the nerve bundles in a heterogeneous punctate pattern. Interestingly, Piezo1 demonstrated a banding staining pattern perpendicular to the nerve evident at the lamina cribrosa (white arrow), where astrocytes are situated (Fig. 4D). Glial fibrillary acidic protein (GFAP) staining was then used to identify astrocyte ^62^ together with Piezo1 at the optic nerve head, where overlapping of both antigens were evident along the glial laminar (Fig. 4E).

### Immunohistochemical identification of Piezo2 in neural retina and optic nerve head of mouse

The expression and distribution of Piezo2 was also examined in the mouse. Piezo2 displayed prominent staining in the nerve fiber layer, showing strong colocalization with beta-tubulin III (Fig. 5A). Piezo2 and beta-tubulin III co-staining continued through the optic nerve head region and into the optic nerve, with the striated staining suggesting localization within the axons. Staining was also detected in RGC soma, with some cells also displaying Piezo2 in beta-tubulin III-positive dendrites. Piezo2-stained cells were found to surround BRN3A positive nuclei (Fig. 5B), confirming Piezo2 expression in RGCs. Minimal staining for Piezo2 was observed in the outer retina.

**Figure 5.**
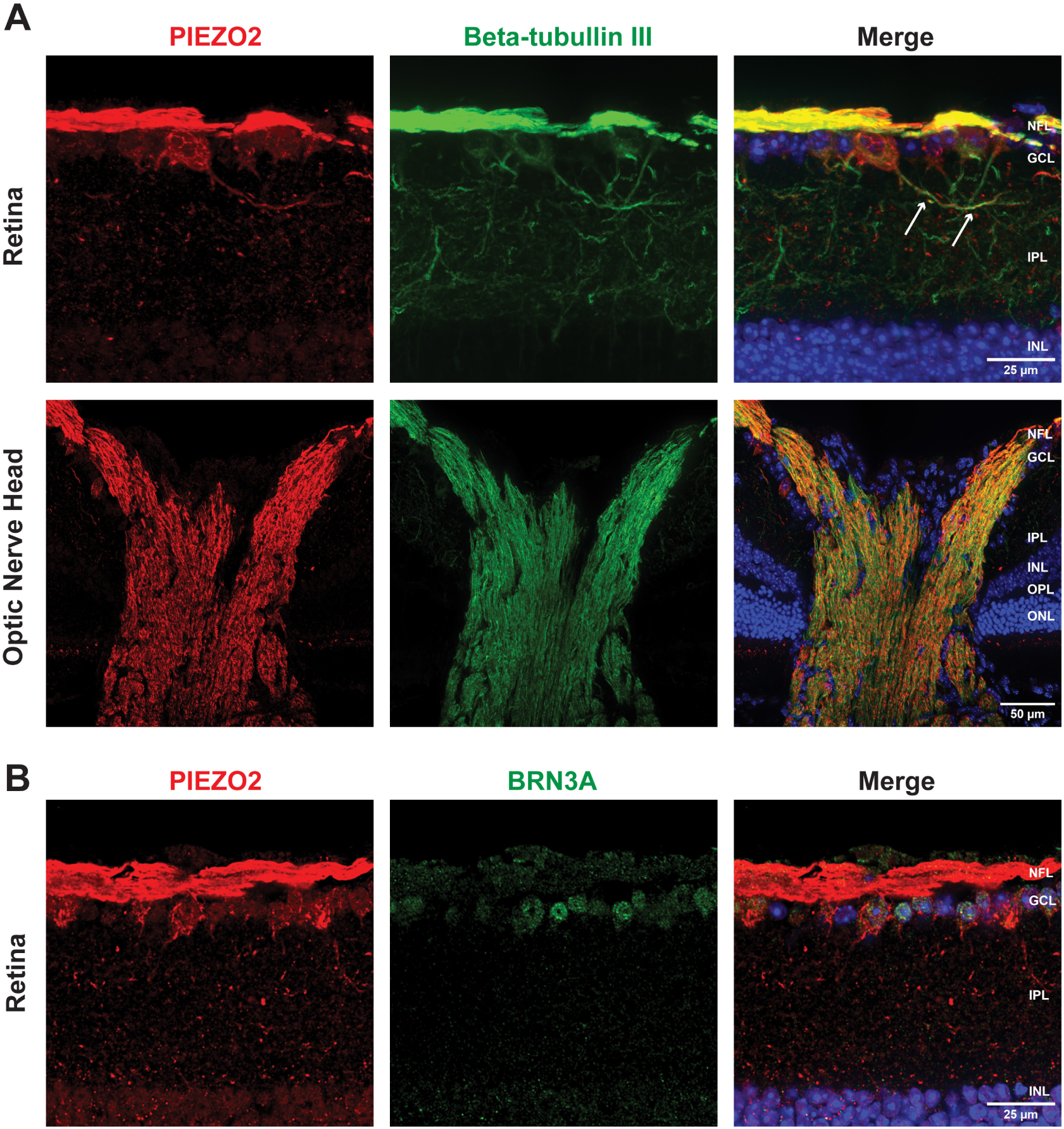
Piezo2 channel immunoreactivity in the mouse retina indicates strong expression in RGC axons: **(A)** Robust expression of Piezo2 in the retina in NFL and GCL. Double labeling with anti-Piezo2 and anti-beta-tubulin III antibodies shows strong colocalization from the soma and axons of RGCs in C57BL/6J eyes continued to the optic nerve after exiting the globe, with banding patterns evident. The staining pattern of the RGCs, especially dendrites and axons, is worth mentioning as a sharper characteristic compared to Piezo1 (white arrow). The inner nuclear layer displayed less staining for Piezo2 than for Piezo1. **(B)** Representative images of double immunolabeling Piezo2 and BRN3A in the retina demonstrated a shared localization of both proteins in the RGCs. Insets show magnified images of the regions enclosed by the white dashed boxes. Images representative of staining in 3 mice.

### Piezo1 and Piezo2 expression in human iPSC-RGCs

To further characterize the expression of Piezo channels and increase the relevance to humans, we examined human induced pluripotent stem cell-derived retinal ganglion cells (iPSC-RGCs). Cells derived using this process are electrically active and express key RGC markers ^29, 63^. Identification was confirmed with iPSC-RGCs staining for markers BRN3A and RBPMS (Fig. 6A). Particulate staining for Piezo1 was detected in the soma and neurites of iPSC-RGCs (Fig. 6B); staining for Piezo2 was similar but more robust. Strong co-localization between Piezo2 and beta-tubulin III was evident, particularly in the neurites (Fig. 6C).

**Figure 6.**
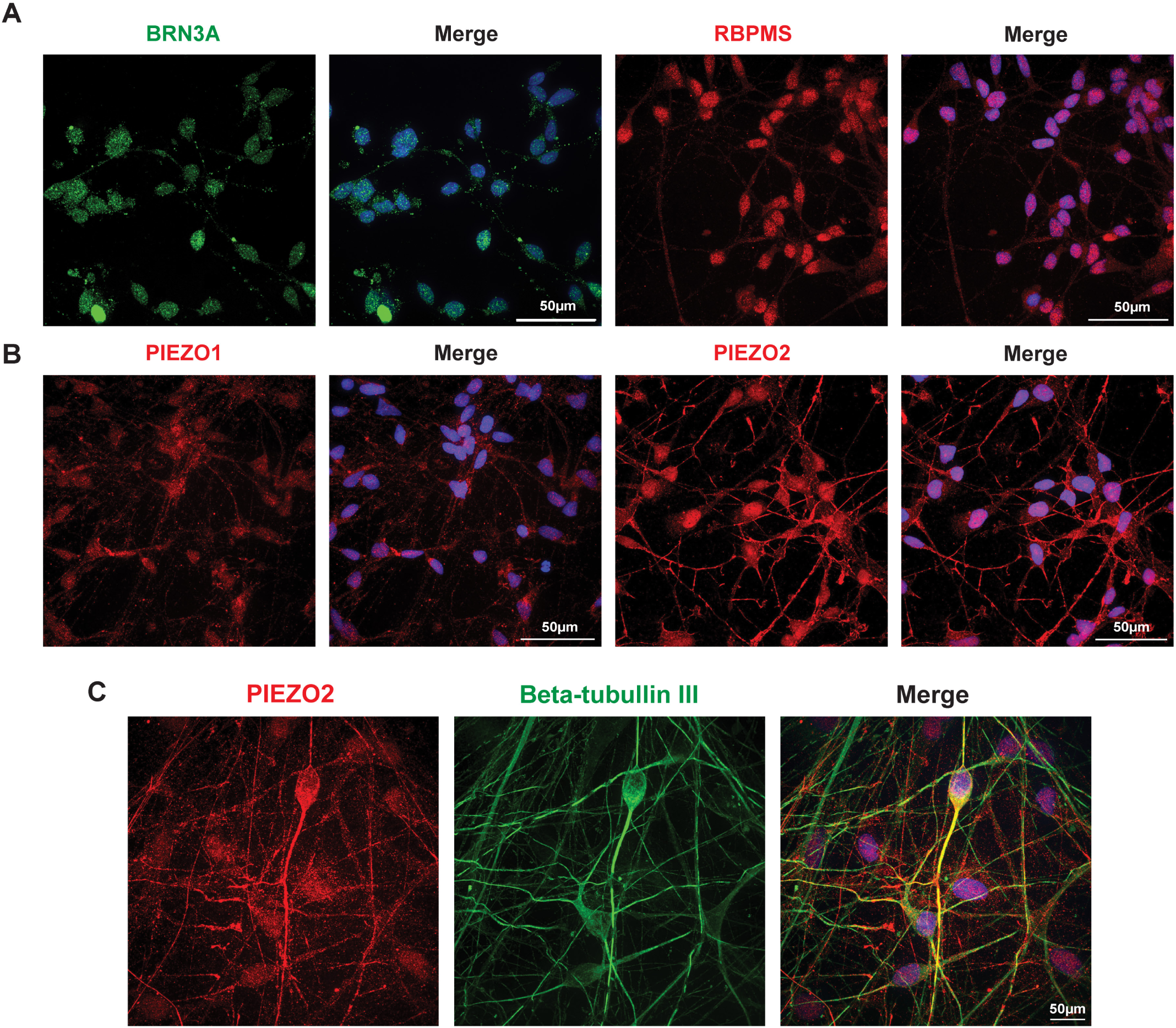
Expression of Piezo channels in iPSC-RGCs: **(A)** iPSC-RGCs were stained for pan retinal ganglion cell markers, Brn3A, and RBPMS to confirm the characterization. Scale bar, 50 μm. **(B)** Representative images showing positive staining of Piezo1 and Piezo2 in iPSC-RGCs confirm data from murine retinal tissues. Scale bar, 50 μm. **(C)** Representative images showing double immunostaining for Piezo2 and beta-tubulin III in iPSC-RGCs. Strong co-localization can be seen from the soma projecting toward the neurite of the cells (n=3; 2 biological replicates).

### Immunohistochemical identification of Piezo1 and Piezo2 in neural retina and optic nerve of rat

Immunolocalized experiments were repeated in the rat retina to support the identification in mouse and human material. Staining of Piezo1 in rat retina was similar to that in the mouse, although with more intense staining observed along the internal limiting membrane (Fig. 7A). Colocalization with beta-tubulin III supported Piezo1 in some RGC soma. Light particulate immunoreactivity of Piezo1 between strands of beta-tubulin III-positive regions was noticed throughout the optic nerve head. Piezo2 staining was also similar in rat to that observed in mouse, with clear colocalization in the nerve fiber layer and weaker staining in the soma and dendrites of RGCs (Fig. 7B). Staining for Piezo2 and beta-tubulin III was also present throughout the optic nerve head of rat tissue.

**Figure 7.**
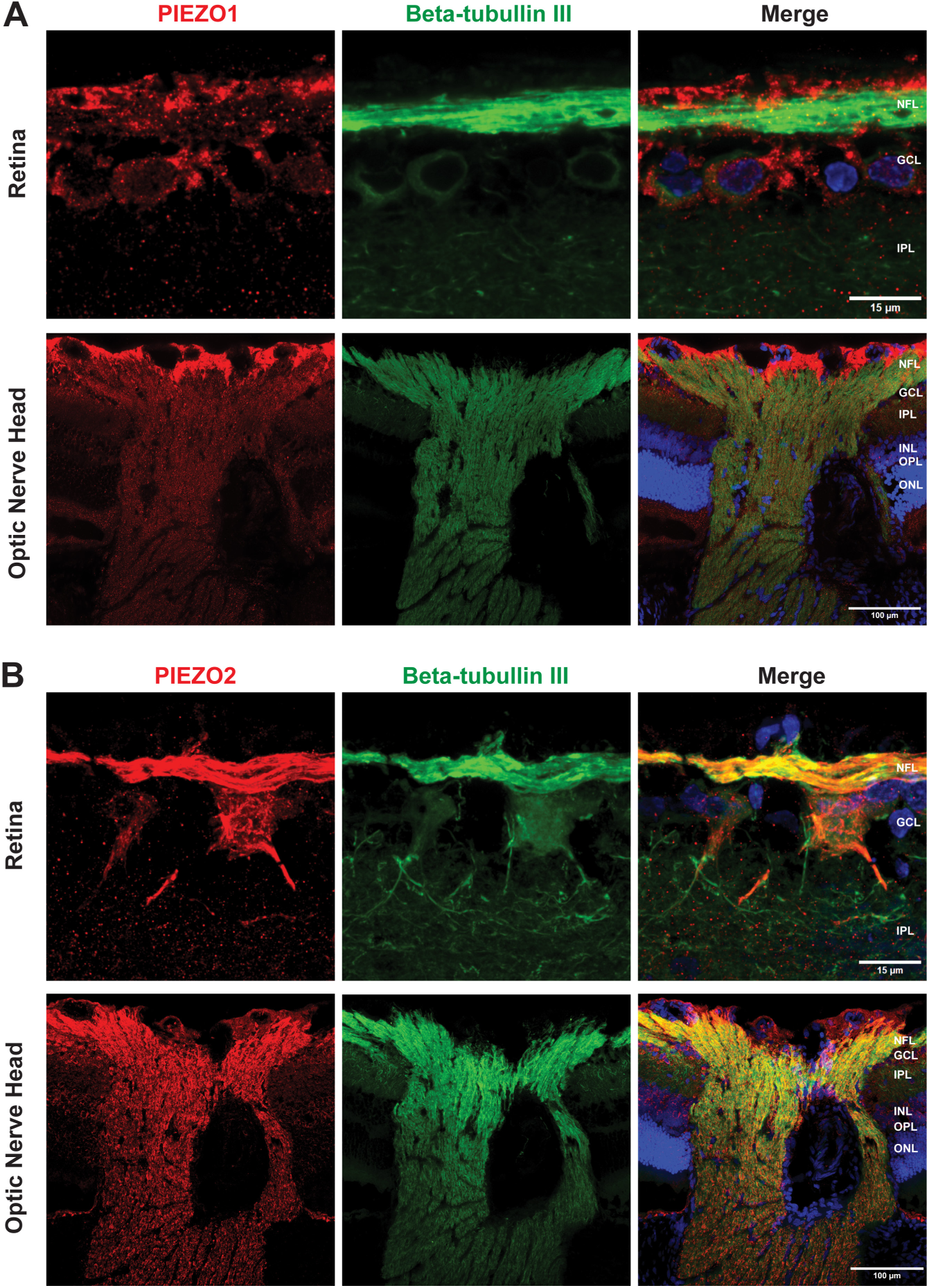
Long Evans rat retina shows similar immunoreactivity for Piezo1 and Piezo2 channels: **(A)** Reciprocal expression of Piezo1 to C57BL/6J eyes was recognized in the retina in NFL, GCL, and sparsely in INL. Double labeling with anti-Piezo1 and anti-beta-tubulin III antibodies show positive staining in RGCs’ soma but not as strong as the end feet of Muller glial cells in the internal limiting membrane. Minimal staining at the RGC dendrites was also noted. Positive staining can also be seen in the ONH area. **(B)** Vigorous expression of Piezo2 in the retina in NFL and GCL. Co-staining with anti-Piezo2 and anti-beta-tubulin III antibodies shows strong colocalization from the soma and axons of RGCs, similarly as in C57BL/6J eyes continued to the optic nerve after exiting the globe. Pronounced expression at the dendrite of the RGCs was also presented. Insets show magnified images of the regions enclosed by the white dashed boxes. Images representative of staining in 3 rats.

## Discussion

The present study provides an analysis of Piezo channels in the retina, using multiple approaches and models to strengthen conclusions. The presence of both Piezo1 and Piezo2 channels was particularly robust in RGCs. The co-localization between beta-tubulin III and Piezo1 suggests channel expression in somas and axons of nerve fibers. The staining of iPSC-RGCs showed a similar pattern, while Yoda1 increased Ca^2+^ in the soma and neurites of cultured mouse RGCs. Robust staining of Piezo 2 was detected in mouse, rat and human RGCs, with particularly strong staining in the axons of the nerve fiber layer and through the optic nerve head. Staining for Piezo1 was also colocalized with markers for astrocytes in the optic nerve head and Müller cells at their end feet. Together, the results implicate Piezo1 and 2 channels in homeostatic processes of adult RGC axons, with contributions suggested from Müller cells and optic nerve head astrocytes.

Analysis of mRNA levels also suggests expression of both *Piezo1* and *Piezo2* in the retina, optic nerve, and the RPE/choroid. While expression of both channels was relatively low in the retina, this likely reflects the presence of the channels in only a few of the retinal cell types. The robust expression of message for *Piezo1* and *Piezo2* in the RPE/choroid was unexpected, given the tissues are non-excitable and not traditionally thought of as mechanosensitive. While immunostaining for both Piezo1 and Piezo2 in ARPE-19 cells supports this identification, confirmation in vivo is currently being investigated.

Dysfunction in neuronal signaling pathways, including RGC function, has been associated with alterations in IOP dynamics ^64^. Early compensatory response in RGCs characterized by increased excitability levels in both ON and OFF subtypes, irrespective of the extent of dendritic pruning, have been identified as a notable feature of the initial stages of neurodegeneration in glaucoma models. Risner et al. observed that following two weeks of microbead-induced IOP elevations in mice, representing an early time-point in disease propagation, there was a noticeable increase in depolarization of the resting membrane potential of RGCs, leading to a lowered excitation threshold. Furthermore, responses to the onset of light stimuli also exhibited increased magnitude, duration, and frequency. However, these enhancements were transient and undetectable at four weeks of sustained IOP elevation ^65^. Tao et al. demonstrated that alterations in RGC function, including an increase in spontaneous firing rate and excitability, were evident only under conditions of microbead-induced IOP elevation at lower magnitudes but not at higher IOP levels ^66^. Strikingly, our findings suggested that Piezo1 may play a role in this early adaptive role in glaucoma as the results emulate what had been described previously *in vivo*, where RGCs modify their physiology to maintain functionality, mitigating affected RGCs to improve signaling between neurons.

The presence of both Piezo1 and Piezo2 in RGC stresses the importance of mechanosensitivity to these neurons. While the two Piezo isoforms are often expressed in different cell types, evidence for the expression of both isoforms is emerging ^11, 67–69^. Simultaneous labeling for Piezo1 and Piezo2 was hampered by a shared host for reliable antibodies, but the expression of both channels in most RGC axons supports co-expression in at least some cells. The co-expression of message for both Piezo1 and 2 was recently described in cells of the anterior eye ^27^. The expression of both Piezo1 and 2 in these neurons is of interest given that they can show synergistic actions ^67^ and may compensate for changes in one another’s expression ^11^. It remains to be determined whether a similar interaction occurs between Piezo1 and Piezo2 in RGC axons. However, the robust expression of both channels in the axons may reflect the importance of the contribution of Piezo channels to the neurons.

### Strengths of the study and comparison to others

Confidence in the findings of the current study is enhanced by several aspects of the experimental design. The molecular and protein data converge, with expression of mRNA found with PCR and qPCR (Fig. 1) in agreement with that found on the protein level with immunohistochemistry (Fig. 4-7). The agreement between expression patterns in the mouse (Fig. 4-5) and rat (Fig. 7) retina, and in human iPSC-RGCs (Fig. 6) also supports the conclusions. The ability of Yoda1 to induce Ca^2+^ influx into RGCs adds a functional confirmation of Piezo1 in RGCs and optic nerve head astrocytes (Fig. 3). Although the specificity of some of the antibodies for Piezo channels used previously was questionable, antibodies with improved characteristics have been developed in the past few years. Our data indicated that knockdown of Piezo channels with siRNA dramatically reduced staining and strengthened support for these antibodies (Fig. S3). Additional support from others validates the staining of the antibodies used in this study, including KO mice models, reporter mice, pre-absorption assay, or genetically modified cell lines ^50–57^.

Investigating the early reactions of RGCs and their microenvironment to initial stressors is pivotal for understanding the pathogenesis of glaucoma. To our knowledge, this is the first study to identify both baseline and scotopic response changes with Piezo1 activation in the retina, using MEA recording. Increases in excitability drive greater energy demand and can lead to a greater accumulation of metabolic by-products, including reactive oxygen species, from mitochondrial ATP production trying to mitigate the energy imbalance of the RGCs, which in turn may lead to RGC dysfunction ^70^. Similar emergence of spontaneous hyperactivity in RGCs has also been detected in animal models of retinal degeneration as the disease progress ^71–74^. Speculations concerning the impact of spontaneous hyperactivity among RGCs on patients’ vision suggest an increase in background noise, potentially disrupting the transmission of normal signals essential for visual perception when processed by various brain circuits. Ultimately, this could lead to a reduced signal-to-noise ratio, characterized by a higher rate of action potentials discharged by ganglion cells ^75, 76^. Piezo1, a mechanically activated ion channel, has been implicated in various physiological processes, including cellular mechanotransduction. In the context of glaucoma, where increased intraocular pressure can lead to mechanical stress on RGCs, Piezo1 could be a critical factor in how these cells adapt to preserve their function or mitigate stress and could potentially open up new avenues for therapeutic strategies targeting RGC protection and neuroprotection, which may complement IOP-lowering treatments and improve outcomes for patients with glaucoma. With no available specific Piezo2 agonist, its involvement cannot be ruled out, as they may have overlapping functions.

The observations above contribute to the understanding of Piezo1 and 2 channels in the posterior eye, confirming some, but not all, previous findings. For example, staining for Piezo2 was quite evenly distributed throughout the retina in an earlier study on mice, although Piezo1 showed some concentration in the ganglion cell layer ^24^. In the guinea pig retina, staining of Piezo1 is also expressed throughout the retina ^25^. A recent study using fluorescent in situ hybridization to overcome issues with earlier antibodies found mRNA for *Piezo1* in selected regions of the mouse ganglion cell layer and inner nuclear layer, although expression of *Piezo2* message was more restricted ^27^. This difference is not unexpected, however, as the highest antigen signal in our study was from the axons where mRNA levels are likely quite low. RNA messages for *Piezo1* and *Piezo2* were previously reported in mouse optic nerve head tissue and in 22% and 43%, respectively, of isolated optic nerve head astrocytes ^77^; while our immunostaining suggests a higher level of expression in cultured optic nerve head astrocytes, this may reflect differences in culture protocols. In a more recent study, Yoda1 increased fibronectin and Ca^2+^ levels in mouse optic nerve head astrocytes with Piezo1, but not Piezo2, localized to optic nerve head astrocytes and increased by stretch ^78^, although this study was exclusively in vitro. Addition of Yoda1 and increased hydrostatic pressure-activated currents and Ca^2+^ transients in retinal capillary cells ^26^; although we did not notice any staining in retinal capillary cells, their detection is best done in wholemounts, and we largely used sections, so cannot rule them out.

### Implications for pathology, neuron growth, and interactions with other mechanosensitive channels

The identification of both Piezo1 and Piezo2 on axons of RGCs as they pass through optic nerve head has direct relevance in glaucoma, as elevated IOP distorts lamina cribrosa of the optic nerve head, causing damage to the axons ^79^, and axonal neuropathy of RGCs is an early sign of glaucomatous damage ^80, 81^. While the role of Piezo channels in this mechanosensitive damage is unclear, their substantial presence in healthy axons suggests a protective contribution under homeostatic conditions. Excessive stimulation may be damaging, however, and the sustained influx of Ca^2+^ is pathological to neurons. Aberrant Ca^2+^ signaling recently indicated dysfunctional cortical signaling after mild traumatic brain injury ^82^, and can disrupt axonal transport ^83^. As mitochondrial dysfunction plays a central role in RGC loss ^84^, recent reports of the activation of mitochondria by Piezo1 may have relevance ^85^.

In addition, the mechanosensitive release of ATP through pannexin hemichannels is implicated after stretch and in eyes with elevated IOP ^86–88^, and stimulation of the P2X7 receptor for ATP can activate microglial cells and injure neurons ^41, 89^. Piezo channels can trigger ATP release from red blood cells ^90^ and interact with pannexin channels in cholangiocytes ^91^, although a Piezo-pannexin-ATP link in the optic nerve head remains to be determined. Recent evidence that repeated elevations in IOP can lead to inflammation following activation of the pannexin1 channel supports this link ^92^. The inflammatory responses accompanying the elevation of IOP are substantial, suggesting a possible upstream role for Piezo channels ^93, 94^.

Elevation of both *Piezo1* and *Piezo2* after stretch *in vivo* may have implications for pathology. Mechanical forces influence expression of Piezo channels; expression of Piezo1 channels in stem cells was increased with substrate stiffness ^95^, while Piezo1 levels in myofibroblasts were increased with 18 or 24 hours of 10% stretch ^96^. In the optic nerve head, expression of Piezo2, but not Piezo1, was upregulated in the DBA/2J mouse model of glaucoma at 10 ^77^ or 15 months ^24^, although the strain was relatively continuous in this model. A two-hour 10% stretch doubled Piezo1 expression in mouse optic nerve head astrocytes ^78^. An ischemia-perfusion model in which IOP was raised to 100 mmHg increased Piezo2 expression at 3 and 5 days, with a return to baseline at 7 days ^24^, while elevating IOP to 30 mmHg for 1 hr did not alter Piezo1 or 2 expression ^77^. The sustained rise in Piezo expression detected in this study suggests the CEI model of ocular hypertension, where IOP is increased to 60 mmHg to generate a non-ischemic pressure rise, produces a sufficient tissue stretch to upregulate Piezo expression. Similar upregulation of specialized excitatory channels, NaV1.6 ^65^ and Trpv1 ^97^,has also been reported, accompanying a chronic rise in IOP. The ability of a single episode of tissue stretch to induce a prolonged rise in Piezo channel expression suggests a mechanism where neural tissue becomes increasingly sensitive to mechanical stain with repeated stimulation. This has implications for brains exposed to repeated traumatic injury, and for eyes exposed to multiple spikes in IOP, as an increase in Piezo expression could increase the Ca^2+^ entry, with negative consequences for neurons.

While Piezo1 has been implicated in the growth of RGC axons, the nature of this influence remains to be determined as inhibition of mechanosentitive channels with GsMTx4 and knockdown of Piezo1 reduced RGC axon growth in Xenopus ^98^, but GsMTx4 increased growth of mouse RGC neurites in culture and Piezo1 agonist Yoda1 reduced their growth ^24^. The role of Piezo channels in the retina may vary with age, as Piezo1 levels were higher in the optic nerve head of neonatal mice than in 3-month-old mice; cultured astrocytes with Piezo1 knocked down showed decreased expression of cytoskeletal and cell cycle genes, suggesting a role in development ^99^. The results from cultured cells above may reflect developmental influences on the cells. Regardless, the presence of Piezo1 on RGC axons in mice 3-6 months of age implies the channel has additional contributions to the homeostatic functions of adult RGCs.

In addition to its own mechanosensitive activity, Piezo channels may also influence the activity of other channels. Piezo channel opening contributes to the mechanosensitive current through TREK-1 channels ^100^, while Piezo1 was identified as the upstream mechanosensor that connects shear stress to TRPV4 opening in endothelial cells ^101, 102^. This could help the transient opening of the Piezo channel pore to drive a more sustained response, particularly at higher levels of strain ^103^. Similar interactions between Piezo channels and TRPV4 would have relevance in the retina, where TRPV4 has long been implicated in mechanosensitive responses ^104–106^. Müller cells showed strong expression of mRNA for both Piezo1 and TRPV4, and the mechanosensitive elevation in Ca^2+^ was only partially reduced in TRPV4^-/-^ mice, suggesting Piezo1 channels may interact with TRPV4 in the mechanosensitive responses of Müller cells ^107^. The Ca^2+^ elevation induced by mechanical strain initiated in the end feet region; while the colocalization of Piezo1 with glutamine synthetase on Müller cell end feet above supports an interaction, more research is needed.

## Conclusions

In summary, this study has identified both Piezo1 and Piezo2 channels in mouse, rat and in human iPSC-RGCs, with expression in soma and axons. Functional assays confirmed the presence of Piezo1 in the soma and neurites of the neurons. Activation of the Piezo1 channel in the explant retina increased RGC excitability at both baseline and scotopic conditions. Sustained elevation of *Piezo1* and *Piezo2* occurred after a single transient stretch. The presence of both Piezo1 and Piezo2 on the axons and soma, and the sustained rise in Piezo expression after a single exposure to stretch may provide a mechanism for enhanced neural sensitivity to repeated insults in pathologies like glaucoma or traumatic brain injury.

## Supporting information

Extended method for supplemental figures

## Acknowledgements

The authors would like to thank Hydar Ali for his generous access to his Varioskan device. This study was supported by R01 EY015537, R01 EY015537S1, R01 EY013434, P30 EY001583, R01 EY023557, Research to Prevent Blindness Unrestricted Grant, Penn Health-Tech Seed Funding.

## Conflict of Interest

All the authors have read the manuscript and declare no conflict of interest.

## Author contributions

PS made substantial contributions to the conception and design of the work, the acquisition, analysis and interpretation of data, and drafted and edited the work; LS made substantial contributions to the conception and design of the work, and the acquisition; SN made substantial contributions to acquisition, analysis and interpretation of data; VRMC made substantial contributions to the acquisition of data; VV made substantial contributions to the acquisition of data; JH made substantial contributions to the acquisition of data, JMOB made substantial contributions to the acquisition of data; JX made substantial contributions to the acquisition of data, analysis and interpretation of data; WL made substantial contributions to the conception and design of the work, the acquisition, analysis and interpretation of data; CHM made substantial contributions to the conception and design of the work, analysis and interpretation of data, and drafted and edited the work. All authors have approved the submitted version and agreed both to be personally accountable for the author’s own contributions and to ensure that questions related to the accuracy or integrity of any part of the work, are appropriately investigated, resolved, and the resolution documented in the literature.

## Data Availability statement

All data generated or analyzed during this study are included in this manuscript and its supplementary information files, or are available from the corresponding author upon reasonable request.

## Notes

### Competing Interest Statement

The authors have declared no competing interest.

