## Extended method for supplemental figures for "Piezo1 and Piezo2 channels in retinal ganglion cells and the impact of Piezo1 stimulation on light-dependent neural activity"

### Supplemental Methods

*Knockdown of Piezo channels with RNA Interference:* ARPE-19 cells were obtained from the American Type Culture Collection (ATCC, Manassas, VA) and grown in a 1:1 mixture of DMEM/F12 with 3 mM L-Glutamine, 100 µg/mL streptomycin, 100 U/mL penicillin, (all obtained from Invitrogen) in the presence of 10% FBS (Sigma, St. Louis, MO) as described [42]. ARPE-19 cells were transfected with 15 nM SMARTpool ON-TARGETplus human Piezo1 siRNA (L-020870-03-0005; Dharmacon), SMARTpool ON-TARGETplus human Piezo2 siRNA (L-013925-02-0005; Dharmacon), or Silencer™ Select Negative Control No. 1 siRNA (4390843; Thermo) using Lipofectamine 3000 (Life Technologies) according to the manufacturer's instructions with media change after 48-hour incubation. The fluorescence intensity was analyzed with ImageJ, with 3 images per each independent sample. The lower threshold was adjusted to remove the background, using the same setting for all images.

*Whole-cell voltage-clamp recording:* Pipettes with resistance of 5-8 MΩ were filled with pipette solution contained (in mM): 0.1EGTA, 20 KCl, 110K gluconate, 10 HEPES, 2 MgATP, 2 Na2ATP (pH 7.2). The bath solution contained (in mM): 140 NaCl, 5 KCl, 10 HEPES, 2 CaCl2, 5 Glucose (pH 7.4). RGCs were voltage clamped from -100 to +80 mV for 300 ms from a holding potential of -60mV. Once baseline currents had been obtained, negative or positive pressure were applied through the patch pipette by HSPC-1 pressure clamp controller (ALA Scientific Intruments). The whole-cell recording was controlled by HEKA 2.53 software (HEKA Elektronik). The currents were amplified with an EPC 10 USB Patch Clamp Amplifier. The signals were low-pass filtered at 1 kHz and sampled at 5 kHz.

*Current-clamp recording:* Recordings were made by using the whole cell configuration of the patch-clamp technique in current-clamp mode with EPC 10 USB amplifier (HEKA Electronics Inc.). Signals were low-pass filtered at 1 kHz and sampled at 10 kHz. The resistance of pipettes was 5-8 MΩ when filled with electrode solution (in mM): 140 KCl, 2 $MgCl_2$, 0.514 $CaCl_2$, 10 HEPES, 1 EGTA, 2 MgATP, 2 Na2ATP (pH 7.3, osmolarity ~300 mOsm, free [$Ca^{2+}$] = 100 nM, EGTA Calculator: http://maxchelator.stanford.edu/CaMgATPEGTA-NIST.htm). The bath solution contained (in mM): 150 NaCl, 5 KCl, 2 CaCl2, 0.5 $MgCl_2$, 10 HEPES, 5 Glucose (pH7.3, osmolarity ~310 mOsm). Cells were maintained at a holding voltage of -60 mV after forming tight seal. To observe spontaneous spikes of RGCs, no constant currents were injected. RGCs manifesting spontaneous firing patterns at 0 holding current had an average resting potential of -57.8±2.6mV (n=4). For spontaneous spikes, only the cells that had spontaneous action potentials were included in the analysis. Membrane potentials were not corrected for liquid junction potentials. No capacitance compensation was employed. Negative pressure (-30 mmHg) was applied through the patch pipette by HSPC-1 pressure clamp controller. All recordings of spontaneous activity were recorded at room temperature. The experiments were discontinued if the recordings became unacceptable and the data were discarded.

*Patch clamp data processing:* All data are expressed as mean ± standard error of the mean. Data processing was performed using Patchmaster 2.53 (HEKA) and MatLab 11.0 (Mathworks). Graphing and statistical analyses were performed using SigmaPlot 11.2 and SigmaStat 3.5 software (Systat Software, Inc.). Values of $P < 0.05$ were considered statistically significant and determined using Student's t-test.
