## Supplementary material for "Piezo1 and Piezo2 channels in retinal ganglion cells and the impact of Piezo1 stimulation on light-dependent neural activity": Figure S1, Figure S2, Figure S3, Figure S4, and Figure S5

### Supplemental Data

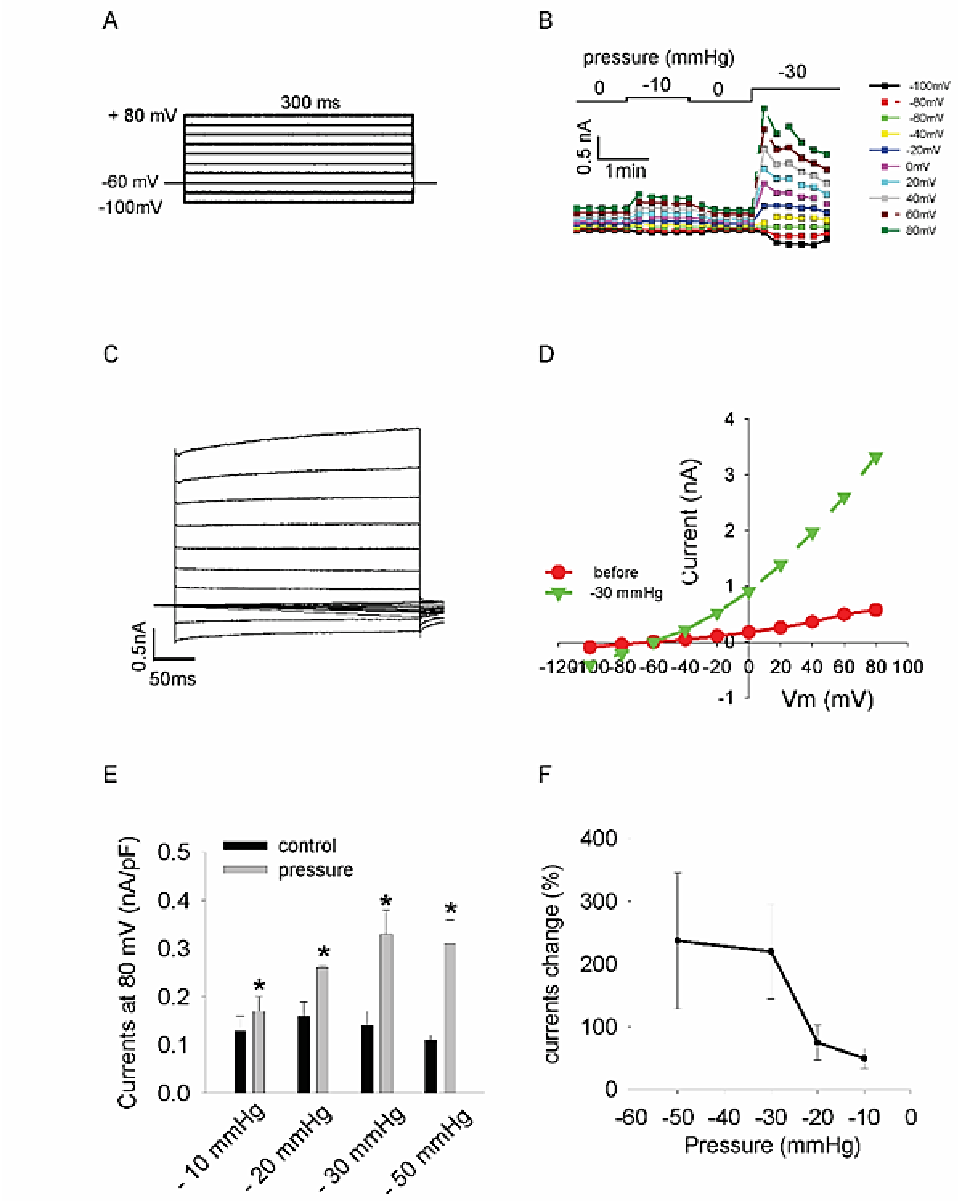


**Figure S1. Negative pressure increased whole cell currents in RGCs. (A)** Voltage protocol used to characterize the whole-cell currents activated by pressure. **(B)** A representative experiment showing the time course of whole-cell currents increase following application of negative pressure. **(C)** A representative whole-cell currents activated by negative pressure (-30mmHg) in a RGC. **(D)** Representative I-V plots before and after application of -30mmHg pressure. **(E)** Overall, applying negative pressure to RGCs led to a significant increase in mean current at +80 mV. n=5 (-10mmHg), n=3 (-20mmHg), n=8 (-30mmHg), n=6 (-50mmHg). *p-value < 0.05. **(F)** Current change percentage at different applied pressure. The cell number tested as in E.

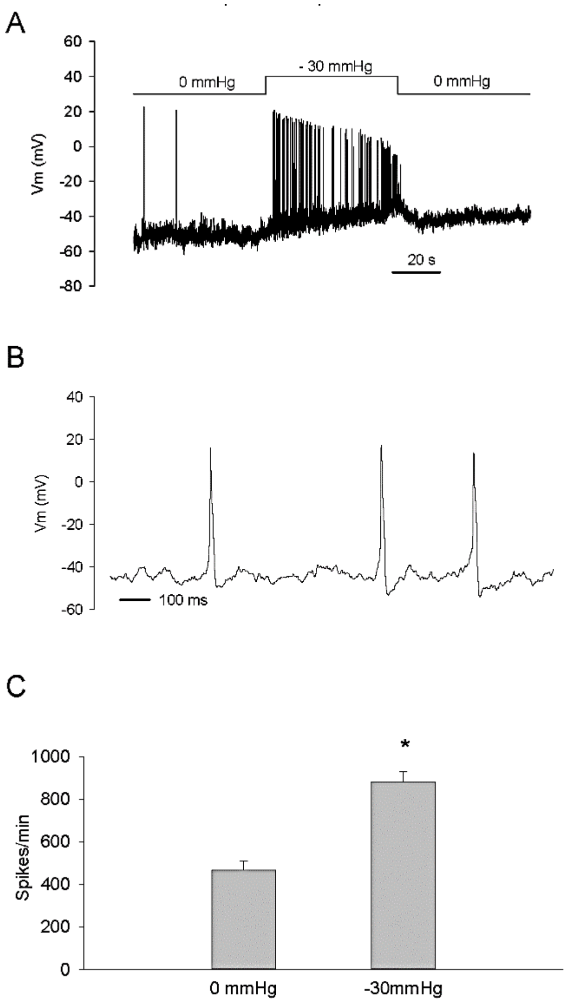


**Figure S2. Pressure increased action potential firing frequency in RGCs. (A)** Representative recording showing pressure initiates increase of action potential firing frequency in a RGC. **(B)** Expanded section from trace in A showing action potential characteristics. **(C)** Average increase of firing frequency (spikes/min). n=4, *p-value < 0.05.

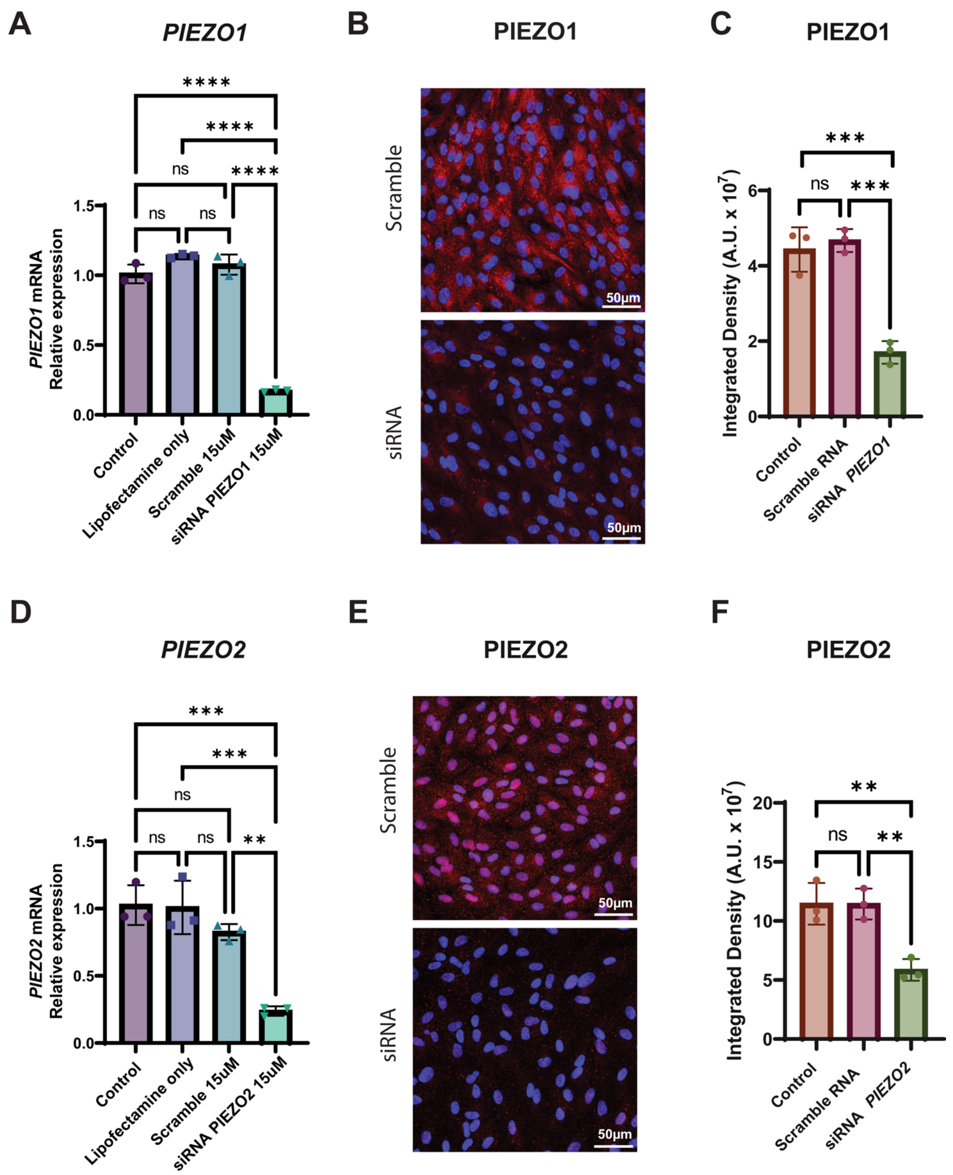


**Figure S3. *Piezo1* and *Piezo2* siRNA knockdown to validate antibody specificity. (A)** qPCR for *Piezo1* expression 48 hours after transfection of ARPE-19 cells with scrambled or *Piezo1* siRNA. **(B)** Representative images of Piezo1 immunoreactivity 72 hours after treatment with scrambled or *Piezo1* siRNA **(C)** Quantitative analysis of fluorescent intensity indicating a significant reduction in cells with knockdown of Piezo1. **(D)** qPCR for *Piezo2* expression 48 hours after transfection with scrambled or *Piezo2* siRNA. **(E)** Representative images of Piezo2 immunoreactivity 72 hours after treatment with scrambled or *Piezo2* siRNA. **(F)** Quantitative analysis Piezo2 immunostaining. (One-way ANOVA tests followed by Tukey's post-hoc test; n = 3). Statistical significance showed at ** p-value < 0.01, *** p-value < 0.001 and **** p-value < 0.0001.

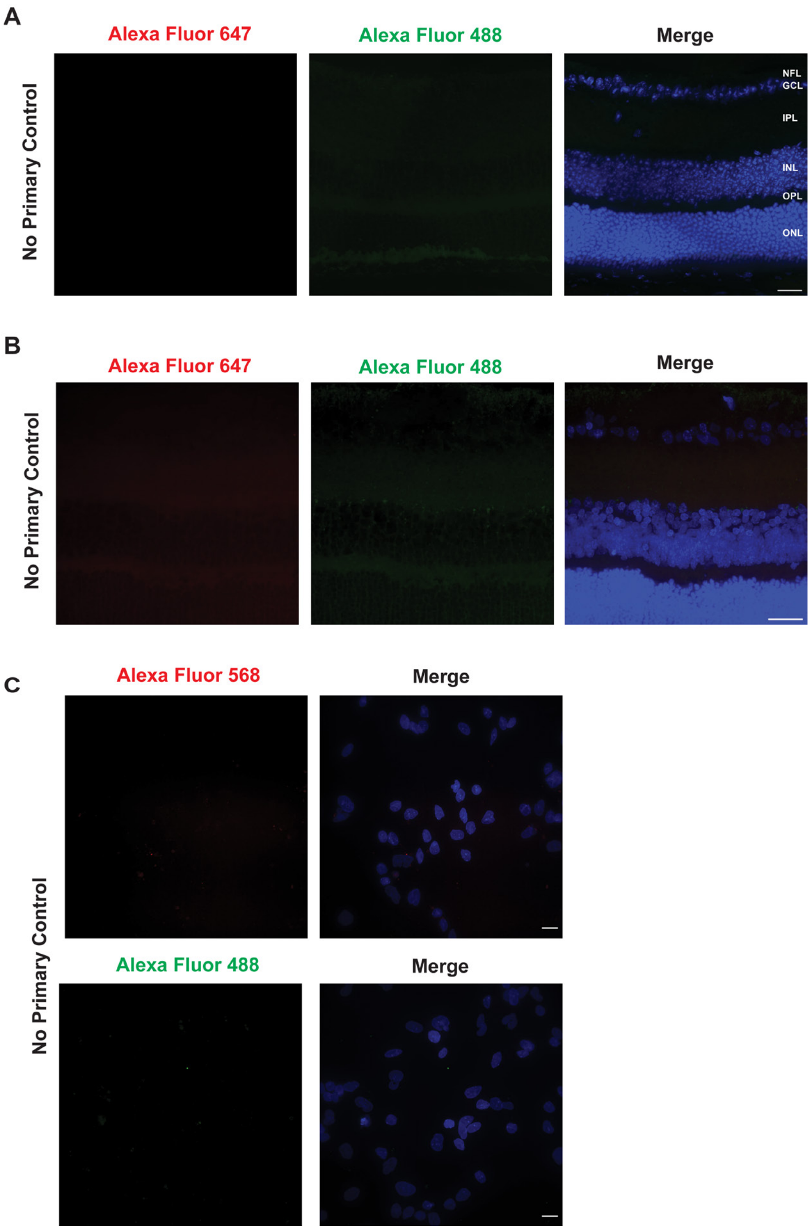


**Figure S4. No primary antibody controls.** (A) Mouse retina. (B) Rat retina. (C) iPSC-RGCs.

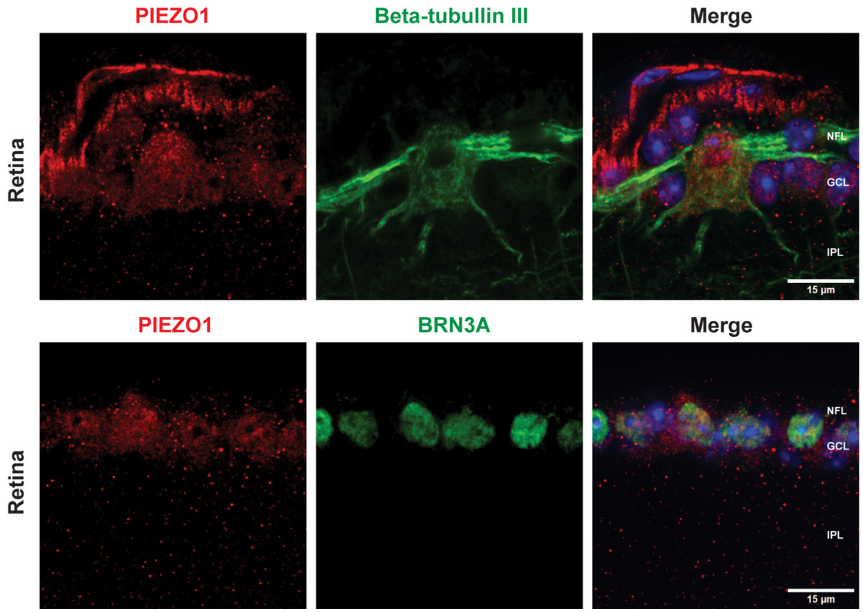


**Figure S5. Second validated anti-Piezo1 antibody also confirmed Piezo1 expression in RGCs.**
